# In mouse and in vitro models, bowel preparation promotes pathogen colonization, translocation and exacerbation of inflammation

**DOI:** 10.1101/2024.11.26.625504

**Authors:** Charlotte A. Clayton, Imogen Porter, Brian D. Deng, Giselle McCallum, Apsara Srinivas, Claire Sie, Jerry Y. He, Alexander Pei, Dominique Tertigas, Deanna M. Pepin, Touran Fardeen, Katharine M. Ng, Sidhartha R. Sinha, Michael G. Surette, Bruce A. Vallance, Carolina Tropini

## Abstract

In the United States an estimated 14 million colonoscopies are performed yearly, each requiring patients to undergo bowel preparation, a laxative cleansing of the intestine’s luminal contents. Despite its widespread use, the effects of bowel preparation on gut physiology and susceptibility to pathogens remains poorly understood, particularly in individuals with compromised gut health. Using mouse and *in vitro* models, we found that bowel preparation with the laxative polyethylene glycol (PEG) rapidly disrupts, transiently increasing susceptibility to infection by *Salmonella* Typhimurium, including a non-motile mutant, and by gut pathobionts derived from ulcerative colitis microbiota. Bowel preparation also facilitated bacterial translocation to extraintestinal sites (mesenteric lymph nodes, liver, and spleen) and exacerbated inflammation in a chemically-induced colitis model. Although these findings are preclinical, they suggest that bowel preparation may have underappreciated risks in vulnerable populations, and warrant further clinical investigation.

## Introduction

In the United States, an estimated 14 million colonoscopies are performed yearly^1^, each requiring patients to undergo bowel preparation (prep), a laxative cleansing of the intestine’s luminal contents. Bowel prep has been shown to cause significant short-term disruptions to the gut microbiota within the first several days post-procedure^2,3^. While bowel prep is widely used and generally considered safe for routine screening colonoscopies, its transient effects on the gut environment, especially in individuals with altered microbiota, remain incompletely understood. Recent studies have shown that colonoscopies have significantly higher rates of infection compared to other screening procedures, with some centers reporting rates of 7-day post-endoscopic infections as high as 132 per 1,000 procedures^4^, and a 9.38-fold increased risk of infection compared to controls that did not undergo these procedures^5^. However, these studies rely on retrospective insurance databases and cannot distinguish whether infections arise from the procedure itself, from interventions such as biopsy, or from preparatory steps like bowel prep. These studies, while limited by retrospective design and confounding variables, suggest a need for further mechanistic investigation into potential transient vulnerabilities. Here, we investigate whether bowel prep alone is sufficient to induce changes in the intestinal environment that could transiently reduce colonization resistance, using reductionist mouse and *in vitro* models to disentangle its specific effects.

Understanding the effects of bowel prep is especially important in vulnerable populations, including patients with inflammatory bowel disease (IBD), comprising both Crohn’s disease (CD) and ulcerative colitis (UC). In healthy individuals, microbiota composition usually returns to normal within days of bowel prep^2,3^. However, the gut microbiota of patients with IBD often harbor microbial species that can act as pathogens under certain conditions. These bacteria, known as pathobionts, are thought to worsen inflammation in IBD^6,7^. Bowel prep may contribute to such disruptions as reports have described IBD exacerbations following colonoscopy^8^ and patients with IBD experience increased risk of post-colonoscopy sepsis and infection^4^. Consistent with these findings, our recent preprint analyzing a national database of quiescent IBD patients undergoing surveillance colonoscopy found an increased likelihood of post-colonoscopy steroid prescriptions, suggesting a risk of delayed symptom exacerbation^9^. Previous studies have shown bowel prep may differentially affect the gut microbiota in patients with or without IBD, leading to long-term changes in microbiota composition several weeks after the procedure^8,10^, as well as adverse effects such as toxic megacolon and increased emergency room visits that have not been mechanistically linked to the procedure itself^11–16^. Understanding how bowel prep affects pathobiont growth and inflammatory state in this population is critical, as patients with IBD undergo more frequent colonoscopies than the general population to monitor disease progression^17^.

In previous work in mice, we showed that long-term (multi-day), low-concentration exposure to the laxative polyethylene glycol (PEG), used as an over the counter laxative and in human bowel prep, compromises the integrity of the intestinal mucus layer^18^, an essential structure for preventing bacterial invasion of the mucosa. Moreover, PEG alters the gut microbiota and selects for bacterial species that can thrive in high osmolality environments^18,19^. Furthermore, we have shown that such high osmolality conditions prevent the growth of commensal bacteria^18^. Importantly, together, these changes have been associated with increased pathogen susceptibility in both mice and humans in some settings^20–22^. However, these prior studies employed low doses of PEG over longer durations than are typical in clinical bowel prep, leaving the specific effects of short-term, high-dose regimens comparatively uncharacterized. Furthermore, despite its routine use, the impact of bowel prep on the gut microbiota, pathogen susceptibility, and its implications in vulnerable populations such as patients with IBD remains insufficiently understood. Specifically, the interactions between bowel prep-induced changes in the gut environment—such as microbiota disruption, mucus depletion, and osmolality changes—and the potential for increased pathogen susceptibility have not been elucidated.

To address this gap, we asked two key questions: (1) How does bowel prep alter the gut microbiota and the intestinal environment, and what are the mechanisms underlying these changes? (2) Does bowel prep worsen host disease state in the context of pathobionts associated with IBD and inflammation? We hypothesize that bowel prep creates an environment in the gut that facilitates the growth and colonization of osmotically resistant pathogens and pathobionts, potentially increasing disease activity.

To answer these questions, in this study we established mouse and *in vitro* models of bowel prep to investigate the effects of acute, high-dose laxative treatment with a time and spatial resolution not achievable in human studies. Specifically, we examined the impacts of bowel prep on the gut microbiota, the local intestinal environment, and host resistance against pathogen colonization by *Salmonella enterica* serovar Typhimurium (*Salmonella* Typhimurium). To robustly colonize the intestines of mice and cause gastroenteritis, this pathogen requires disruption of the microbiota, such as via antibiotic^23,24^ or other drug treatment^25^. We found that bowel prep with PEG transiently altered the gut environment in multiple ways, temporarily increasing its susceptibility to colonization by *Salmonella* Typhimurium, including a non-motile mutant normally unable to invade the gut. In addition, gut preparation promotes the translocation of these bacteria from the gut to extraintestinal organs, such as nearby lymph nodes, the liver, and spleen. In a human IBD microbiota DSS colitis model, bowel prep worsened colitis severity and increased the levels of Enterobacteriaceae pathobionts in extra-intestinal sites, suggesting that bowel prep can transiently lower host defenses in the context of intestinal inflammation. These results highlight bowel prep as a tractable experimental model for dissecting colonization resistance, and suggest that, under certain conditions, it may transiently reduce host defenses in ways that warrant further investigation—particularly in individuals with microbiota rich in pathobionts.

## Results

### Bowel prep disrupts intestinal osmolality, the mucus layer, and short-chain fatty acid levels in the mouse gut

We hypothesized that bowel prep would cause a significant disruption of the gut environment leading to reduced bacterial abundance and depletion of key microbial metabolites, given the severe diarrhea induced by this procedure. To mimic the human bowel prep process, we investigated the impact of short-term, high-concentration PEG exposure on the gut’s mucosal lining. We orally gavaged C57BL/6J mice with PEG (bowel prep) or water (vehicle) four times, at 20-minute intervals (**Fig. 1A**), and assessed changes to the cecal and colon environments 6 hours later. Bowel prep-treated mice had an average cecal osmolality that was 1.7-fold higher than vehicle-treated controls (724 vs. 438 mOsm/Kg, p = 0.00017) (**Fig. 1B**), a higher cecal mass, indicative of osmotic diarrhea, and increased water excretion during bowel prep (**Fig. S1A**). In fixed, stained, and imaged tissue sections, the distal colon of vehicle-treated mice displayed the expected thick, continuous mucus layer (**Fig. 1C**). Conversely, bowel prep-treated mice revealed a largely depleted mucus layer (**Fig. 1C**) and loss of luminal contents including bacteria (**Fig. S1B**) in the colon, despite having similar mass (**Fig. S1A**). Additionally, both the percentage of distal colon epithelium covered by mucus, as detected by UEA-1 staining of fucosylated glycans, and the average mucus thickness, were significantly reduced (**Fig. 1C**). Upstream in the cecum, the mucus layer in vehicle-treated mice was more hydrated and patchier than in the distal colon (**Fig. 1D)**, as expected^26,27^. Here, the percentage coverage and average thickness of UEA-1 stained mucus were also significantly lower (**Fig. 1D)**. Similar trends were observed in WGA-stained mucus, where WGA binds to N-acetylglucosamine and sialic acid residues in glycoproteins (**Fig. S1C**). These findings indicate that even short-term, high-concentration PEG exposure can severely disrupt the protective mucus barrier in the gut.

**Figure 1.**
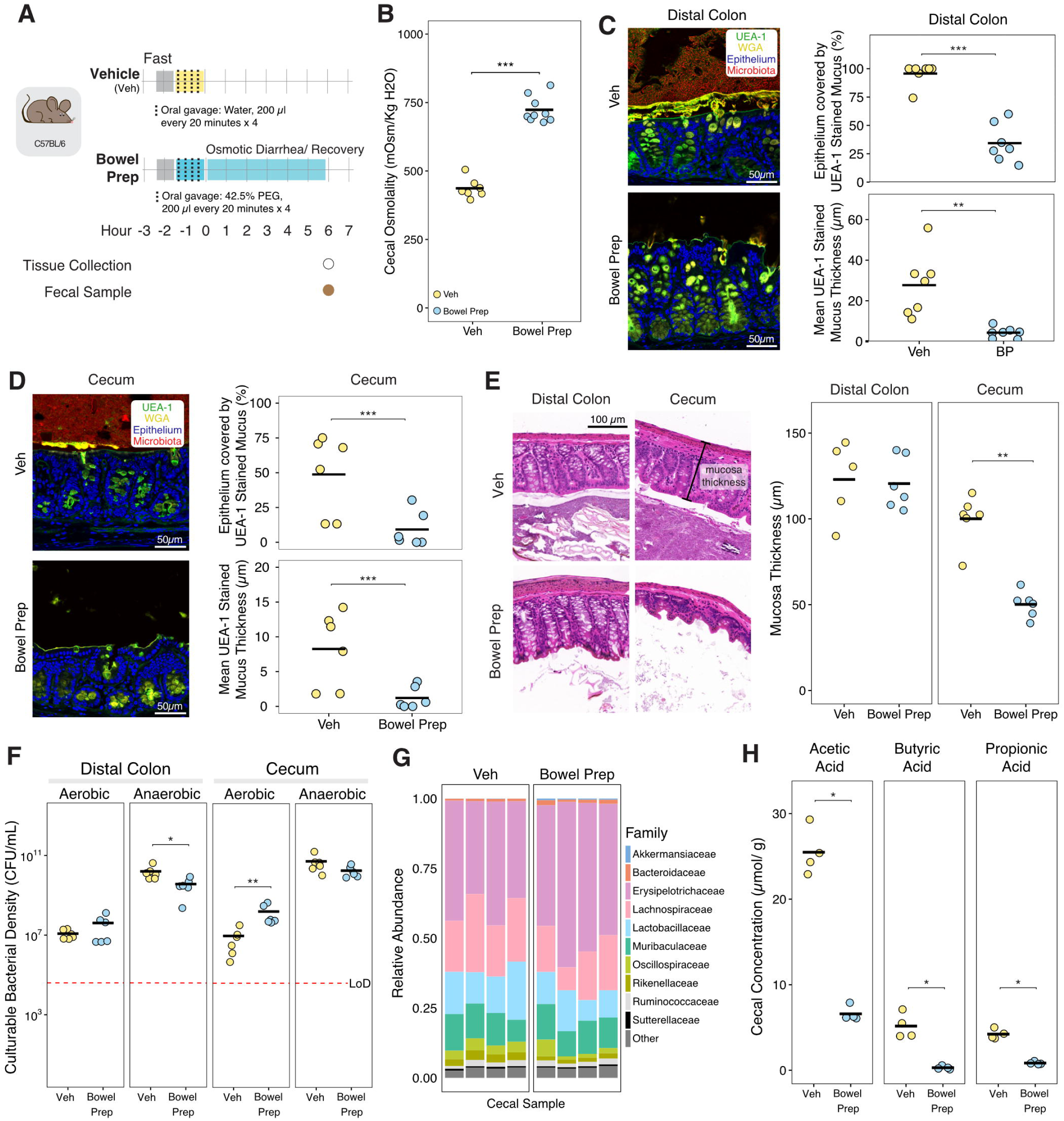
Bowel preparation with a laxative (polyethylene glycol, PEG) disrupts intestinal osmolality, the mucus layer, and short-chain fatty acid (SCFA) levels 6 hours post-procedure in the mouse gut. (A) Schematic representation of our mouse model for bowel preparation (prep). Mice were orally gavaged four times with 42.5% PEG in water (bowel prep) or water (vehicle, Veh), at 20-minute intervals. (B) Cecal osmolality was significantly higher in bowel prep- vs vehicle-treated mice (Veh *n*=7, Bowel Prep *n*=9). (C) *Left*: 6 hours post-treatment, a representative confocal micrograph of the distal colon shows the mucus layer (indicated by staining with UEA-1 [*green*] and WGA [*yellow*]) is considerably thinner in bowel prep- vs vehicle-treated mice. Epithelial cell nuclei (*blue*) were stained with DAPI. *Right*: The percentage of epithelium covered by mucus (*top*) and mucus layer thickness (*bottom*), quantified using UEA-1 fluorescence, were significantly lower in bowel prep- vs vehicle-treated mice (Veh *n*=7, Bowel Prep *n*=6, two independent experiments). (D) *Left*: a representative confocal micrograph of the cecum 6 hours post-treatment, stained and analyzed as in (C). *Right*: The percentage of epithelium covered by mucus (*top*) and mucus layer thickness (*bottom*), quantified using UEA-1 fluorescence (*green*) in bowel prep- vs vehicle-treated mice (Veh *n*=7, Bowel Prep *n*=6, two independent experiments). (E) *Left*: Representative images of hematoxylin and eosin (H&E)-stained sections of the distal colon and cecum 6 hours post-treatment. *Right*: Colon and cecum mucosa thickness in the stained sections (Veh *n*=6, Bowel Prep *n*=6). (F) Loads of culturable anaerobic bacteria 6 hours post-treatment in the distal colon and cecum (Veh *n*=6, Bowel Prep *n*=6). (G) 16S rRNA sequencing of the contents of the cecum 6 hours post-treatment in bowel prep- and vehicle-treated mice. (H) Abundance of three SCFAs in the cecum 6 hours post-treatment, measured using gas chromatography-mass spectrometry (Veh *n*=4, Bowel Prep *n*=4). **Statistics**: For all comparisons, statistical significance was assessed using the Wilcoxon ranked-sum test. p > 0.05; ns (not significant, not shown), p < 0.05; *, p < 0.01; **, p < 0.001; ***, p < 0.0001; ****. **Abbreviation**: LoD, limit of detection.

We wondered whether the disruption of the protective mucus layer might lead to abnormalities in the tissue. We therefore assessed pathology in hematoxylin and eosin (H&E)-stained sections of the cecum and distal colon, including in the lumen, epithelium, mucosa, and submucosa. No immune cell infiltration was observed in samples from bowel prep- or vehicle-treated mice; overall, no difference in tissue pathology was observed between treatment groups (**Fig. S1D)**. However, in the cecum, mucosa thickness was significantly lower in bowel prep- vs vehicle-treated mice (50.2 vs 100.1 µm, p = 0.002; **Fig. 1E**). This reduction in mucosa thickness was not observed in the distal colon (120.6 vs 123.0 µm, p = 0.7922; **Fig. 1E**). This suggests that bowel prep–induced excretion of water, which reduces mucosa thickness, is not uniform throughout the gut.

Next, we measured how bowel prep treatment affects the gut microbiota in mice. We hypothesized that the bacterial loads would be significantly decreased, and the membership changed. To test this hypothesis, we cultured bacteria from the gut both anaerobically and aerobically to measure bacterial loads, as diarrhea is thought to increase oxygen levels in the gut^28^. In the distal colon and small intestine, culturable anaerobic bacterial loads were lower in bowel prep- vs vehicle-treated mice (3.6 x 10^9^ vs 1.6 x 10^10^ CFU/mL, p = 0.01 and 8.4 x 10^8^ vs 1.0 x 10^10^ CFU/mL, p = 0.009, respectively); however, in the cecum bacterial loads did not differ between the two treatment groups (**Fig. 1F, Fig. S1E**). By contrast, culturable aerobic bacterial loads in the distal colon and small intestine – although representing a much smaller fraction of the total counts (0.07% and 0.03% in the vehicle-treated mice, respectively)- did not differ between the two treatment groups. In the cecum, they were 17-fold higher in bowel prep- vs vehicle-treated mice (1.6 x 10^8^ vs 9.6 x 10^6^ CFU/mL, p = 0.005; **Fig. 1F, Fig. S1E**). These findings indicate an overall reduction in microbial load due to bowel prep, and the expansion of aerobic bacteria, consistent with oxygenation of the gut environment. The composition of the cecal microbiota 6 hours post-treatment, as measured by 16S rRNA sequencing, was largely similar in bowel prep- vs vehicle-treated mice (**Fig. 1G, S1H**). Combined with the bacteria load data, this observation suggests that 16S rRNA sequencing is measuring a portion of the microbiota that is either not culturable, or no longer alive post-bowel prep.

Given the changes in microbial load, we expected to find lower levels of microbial metabolites in the gut. To test this hypothesis, we measured individual short-chain fatty acid (SCFA) levels in the cecum’s contents, as major site of microbial activity in the mouse intestine. SCFA levels were significantly lower in bowel prep- vs vehicle-treated mice (3.9-, 17.4- and 5-fold lower, p = 0.03 for acetic, butyric and propionic acid, respectively; **Fig. 1H, S1F**). We also measured the pH of the cecal contents as it is affected by SCFA levels and can impact bacterial growth and composition^19^; however, cecal pH did not significantly differ between the treatment groups (**Fig. S1G**). This suggests that while bowel prep markedly reduces SCFA production, residual levels or buffering by other compounds may be sufficient to maintain luminal pH within a relatively stable range.

Finally, we tested whether bowel prep affected intestinal permeability. We gavaged mice with FITC-Dextran two hours after bowel prep, and collected serum via cardiac punch six hours afer bowel prep. We measured the level of serum FITC fluorescence against a background control, and found no significant difference between mice that received bowel prep or a water vehicle control (**Fig. S1I**).

Altogether, these findings support the hypothesis that bowel prep leads to a significant disruption of the gut environment and impairs microbial metabolism. Given the importance of SCFAs and mucus to enteric pathogen infection^29,30^, these results suggest that the post bowel-prep gut environment may provide diminished colonization resistance against invading bacteria.

### Bowel prep promotes *Salmonella* Typhimurium colonization, translocation, and pathology in mice

Having observed that bowel prep treatment reduces the thickness and coverage of the gut mucus layer, SCFA levels and increases intestinal osmolality (**Fig. 1B**), we hypothesized that enteric pathogens would be able to bypass colonization resistance more easily in bowel prep- vs vehicle-treated mice. Specifically, given that commensal microbiota members are highly sensitive to osmolality^18,19^, we reasoned that an osmotically-resilient pathogen may be advantaged during bowel prep. To test this hypothesis, we measured growth rates of *Salmonella* Typhimurium, an enteric bacterial pathogen known for its ability to withstand osmotic stress in high-salt environments^31^, in growth media adjusted with PEG to various osmolality levels. At an osmolality comparable to that observed in the post-bowel prep gut of mice (∼800 mOsm/Kg), *Salmonella* Typhimurium growth in culture was indistinguishable from growth under normal osmotic conditions (∼450 mOsm/Kg; **Fig. 2A**). Next, we challenged bowel prep- and vehicle-treated mice with a standard infectious dose of 10^6^ CFU *Salmonella* Typhimurium via oral gavage (**Fig. 2B**). Whereas vehicle-treated mice showed no detectable levels of *Salmonella* Typhimurium in their feces post-treatment, bowel prep-treated mice showed high levels of colonization (**Fig. 2C**). One day after inoculation, fecal *Salmonella* Typhimurium levels in bowel prep-treated mice reached 2 x 10^9^ CFU/mL; these levels persisted at least three days post-inoculation, indicating colonization and expansion of the pathogen in the intestinal tract (**Fig. 2C**). This finding was not sex-dependent (**Fig. S2A**). These results support our hypothesis that bowel prep–induced disruption of the gut environment facilitates colonization and growth by osmotically resistant enteric pathogens.

**Figure 2.**
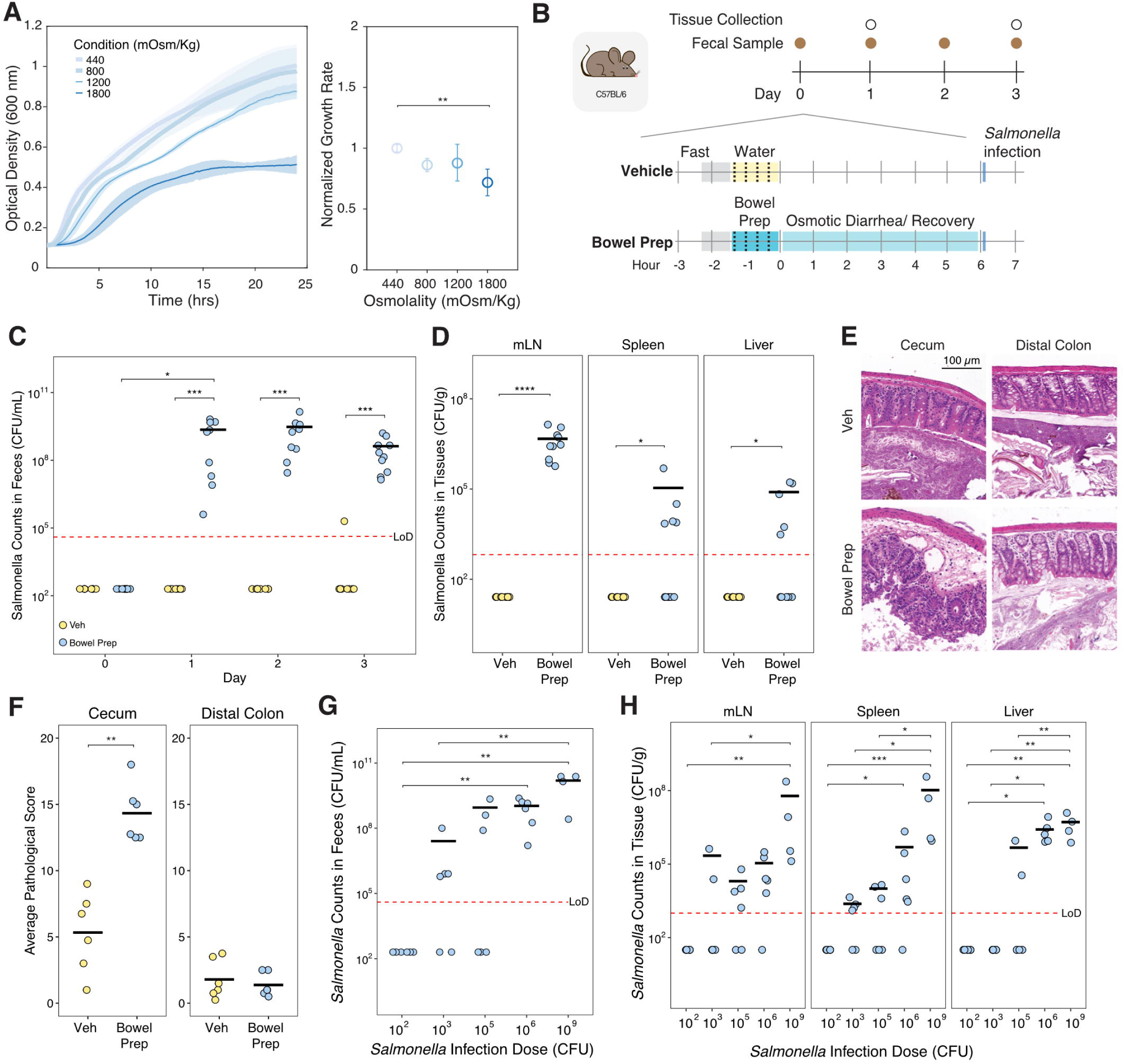
Bowel prep promotes *Salmonella enterica* serovar Typhimurium colonization, translocation, and gut pathology in mice. (A) *Salmonella* Typhimurium growth rates under normal (∼450 mOsm/Kg) and osmotic conditions comparable to those in the post-bowel prep gut of mice (∼800 mOsm/Kg) (each osmolality *n*=4). Growth curves were estimated by measuring optical density (OD600) over time in aerobic media with various osmotic levels (obtained with varying PEG concentrations). Growth rates were compared using a one-way ANOVA with Tukey’s post-hoc test for multiple comparison (each osmolality *n*=4). (B) Schematic of the bowel prep-pathogen mouse model. Mice were infected with *Salmonella* Typhimurium 6 hours after bowel prep with PEG. (C) *Salmonella* Typhimurium counts in the feces of bowel prep- vs vehicle-treated mice (Veh *n*=8, Bowel Prep *n*=10). Differences between timepoints were analyzed with a Wilcoxon ranked-sum test, and differences between bowel prepped-samples at different timepoints were analyzed with a Friedman test followed by Nemenyi post-hoc test. (D) *Salmonella* Typhimurium translocation levels from the gut to the mesenteric lymph nodes (mLN), liver, and spleen three days after inoculation (Veh *n*=8, Bowel Prep *n*=10). Counts in the extraintestinal organs were measured via plating. Differences between groups were analyzed with a Wilcoxon ranked-sum test. (E) Representative images of H&E-stained sections of the distal colon and cecum in bowel prep - vs vehicle-treated mice inoculated with 10^6^ CFU of *Salmonella* Typhimurium. (F) Histopathological scores of the distal colon and cecum sections from (E) (Veh *n*=6, Bowel Prep *n*=6). Differences between groups were analyzed with a Wilcoxon ranked-sum test. (G) Fecal levels of *Salmonella* Typhimurium inoculated with different doses (100-10^6^ CFU *n*=6, 10^9^ CFU *n*=4) 3-days post treatment with bowel prep. Differences between groups were analyzed with Kruskal-Wallis followed by Dunn’s post-hoc test. (H) *Salmonella* Typhimurium translocation levels from the gut to the mesenteric lymph nodes (mLN), spleen, and liver in bowel prep-treated mice 3 days post bowel prep in the mice from (G). Differences between timepoints were analyzed with Kruskal-Wallis followed by Dunn’s post-hoc test. p > 0.05; ns (not significant, not shown), p < 0.05; *, p < 0.01; **, p < 0.001; ***, p < 0.0001; ****. **Abbreviation**: LoD, limit of detection.

Next, we investigated whether *Salmonella* Typhimurium could translocate from the gut to extraintestinal organs. Given the removal of the mucus layer and disruption of the microbiota, we hypothesized that *Salmonella* Typhimurium would be able to reach extra-intestinal organs. Supporting this hypothesis, three days post-inoculation, we identified significant colonization in the mesenteric lymph nodes (4.2 x 10^6^ CFU/g), spleen (4.2 x 10^4^ CFU/g), and liver (3.2 x 10^4^ CFU/g) of bowel prep-treated mice (**Fig. 2D**). No colonization of these organs was observed in vehicle-treated mice, indicating that in normal conditions *Salmonella* Typhimurium translocation is a very rare event.

We also used H&E staining to visualize the host tissue responses to *Salmonella* Typhimurium in the mouse cecum and distal colon and to assess its integrity three days post infection. Given *Salmonella* Typhimurium was able to translocate to extra-intestinal organs, we hypothesized that there would be significant pathology in the intestinal tissues. Supporting this hypothesis, in the cecum, combined histopathological scores for the lumen, epithelium, mucosa, and submucosa were significantly higher in bowel prep- vs vehicle-treated mice (**Fig. 2E,F**). No significant difference in histopathological scores between the treatment groups was found for the distal colon, suggesting that during bowel prep the cecum was the primary site of infection.

We next compared the efficacy of *Salmonella* Typhimurium colonization in bowel prep to the field standard model, which involves pre-treating mice with antibiotics to deplete colonization resistance by native gut microbiota^32^. One day after infection, the maximum fecal burden (as a measure of highest infection level) of *Salmonella* Typhimurium was 11-fold higher in streptomycin- vs bowel prep-treated mice (**Fig. S2B**). This burden persisted for three days after infection (**Fig. S2C**). However, *Salmonella* Typhimurium levels in the extraintestinal organs were similar between the streptomycin- and bowel prep-treated mice (**Fig. 2D, S2D**) and significant pathology was observed in both models at comparable levels (**Fig. 2E,F, S2E,F**). The similar levels of extra-intestinal translocation despite differing intestinal levels suggest that *Salmonella* Typhimurium expansion in the organs, rather than intestinal abundance, may dictate the ultimate organ colonization levels. Altogether, these findings indicate that bowel prep alone, without antibiotics, can facilitate expansion of *Salmonella* Typhimurium in the gut and its translocation to extraintestinal sites.

Finally, we investigated the minimum dose required for *Salmonella* Typhimurium to colonize the gut of mice subjected to bowel prep. We hypothesized that a dose lower than 10^6^ CFU would still enable robust gut colonization and translocation by *Salmonella* Typhimurium, as bowel prep treatment would create a gut environment vulnerable to even low levels of pathogen. Supporting this hypothesis, among bowel prep-treated mice, inoculation with as few as 1,000 CFU led to most mice becoming colonized (**Fig. 2G**). In addition, in bowel prep-treated mice, *Salmonella* Typhimurium translocation levels three days after inoculation largely increased with the size of the dose administered (**Fig. 2H**). These findings indicate that a much lower minimum dose of *Salmonella* Typhimurium is required to colonize the gut after bowel prep, and the higher the dose, the higher the level of translocation to extraintestinal organs.

### Gut resistance to *Salmonella* Typhimurium recovers over time after bowel prep

Considering our finding that bowel prep treatment increased the susceptibility of mice to colonization by *Salmonella* Typhimurium, we investigated which physiological differences might be responsible. Having observed that bowel prep treatment significantly increased pathological scores in the cecum (**Fig. 2E,F**), we characterized the cecal environment in mice at different timepoints post-bowel prep. Cecal osmolality recovered rapidly after bowel prep treatment (765 mOsm/Kg); 24 hours after bowel prep, values were similar to those at baseline (479 vs 423 mOsm/Kg, p = 0.02) (**Fig. 3A**). Cecal mucus coverage and thickness also recovered to baseline values 48 hours after bowel prep (**Fig. 3B, S3A–C**). Furthermore, the cecal and fecal microbiota were depleted by bowel prep, with species diversity levels remaining lower than baseline at 24 and 48 hours after bowel prep treatment before recovering by 72 hours (**Fig. 3C, S3E**). Microbiota composition, as measured by Bray Curtis dissimilarity values, was similarly disrupted following bowel prep (**Fig. S3F**). Finally, SCFA concentrations, particularly for butyrate, remained depleted in bowel prep-treated mice 24 hours after treatment but recovered by 48 hours (**Fig. 3D, S3D**).

**Figure 3.**
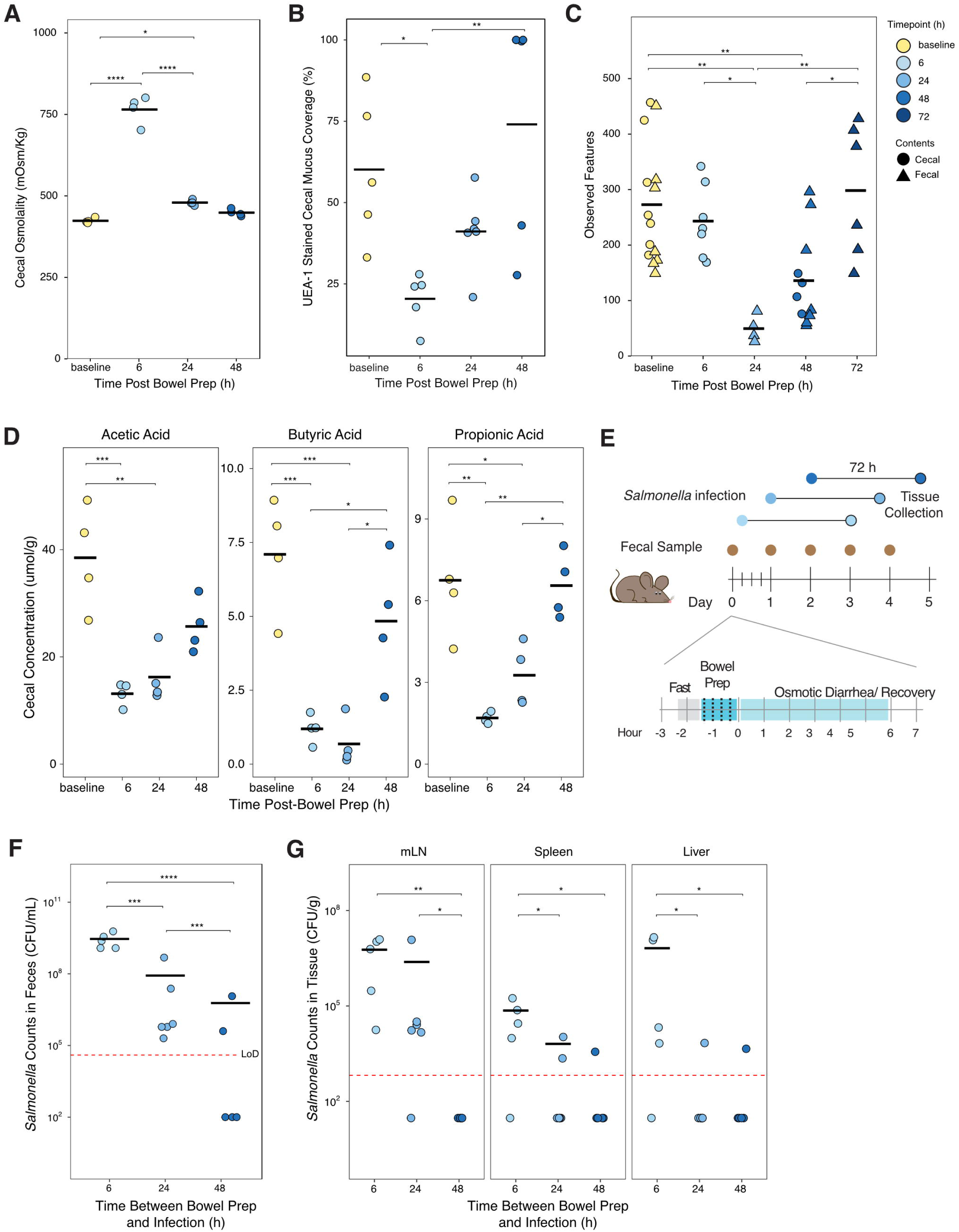
Gut susceptibility to *Salmonella* Typhimurium is reduced by 24 hours after bowel prep. (A) Cecal osmolality at different timepoints after bowel prep with PEG (each timepoint *n*=4). Differences between timepoints were analyzed using t-tests. (B) Percent mucus coverage of the cecal epithelium, as measured with UEA-1 staining (baseline, 6h, 48h *n*=5, 24h n=6, three independent experiments). Differences between timepoints were analyzed using a one-way ANOVA with Tukey’s post-hoc test for multiple comparisons. (C) Number of observed features (alpha diversity) in the cecal and fecal microbiome, determined from 16S rRNA sequencing (baseline *n*=14, 6h *n*=7, 24h *n*=4, 48h *n*=11, 72h *n*=6, four independent experiments). Differences between timepoints were analyzed using a one-way ANOVA with Tukey’s post-hoc test for multiple comparisons. (D) Abundance of three short-chain fatty acids in the cecum, measured by gas chromatography-mass spectrometry (each timepoint *n*=4). Differences between timepoints were analyzed with Kruskal-Wallis followed by Dunn’s post-hoc test. (E) Schematic of bowel prep–pathogen mouse model with *Salmonella* Typhimurium inoculation at 6, 24 and 48 hours after bowel prep treatment. (F) Fecal levels of *Salmonella* Typhimurium (6h *n*=5, 24h *n*=6, 48h *n*=5, two independent experiments) 72 hours post-inoculation. Differences between timepoints were analyzed with Kruskal-Wallis followed by Dunn’s post-hoc test. (G) *Salmonella* Typhimurium translocation from the gut to the mesenteric lymph nodes (mLN), liver, and spleen (6h *n*=5, 24h *n*=6, 48h *n*=5, two independent experiments). Culturing of organs to obtain pathogen counts was performed 72 hours post-inoculation. Differences between timepoints were analyzed with Kruskal-Wallis followed by Dunn’s post-hoc test. p > 0.05; ns (not significant, not shown), p < 0.05; *, p < 0.01; **, p < 0.001; ***, p < 0.0001; ****. **Abbreviation**: LoD, limit of detection.

To investigate whether these changes to the gut environment impact resistance to pathogen colonization, we challenged mice with 10^6^ CFU *Salmonella* Typhimurium 6, 24, and 48 hours after bowel prep treatment (**Fig. 3E**). Seventy-two hours after inoculation, fecal *Salmonella* Typhimurium levels were high in mice inoculated 6 hours after bowel prep treatment but progressively lower when inoculated 24 or 48 hours after treatment (**Fig. 3F**). Similarly, 72 hours after inoculation, *Salmonella* Typhimurium levels in the mesenteric lymph nodes were high in mice inoculated 6 and 24 hours after bowel prep treatment, but lower in mice inoculated 48 hours after treatment (**Fig. 3G**). Together these data suggest that the perturbations caused by bowel prep in mice are transient, and recovery begins within 48 hours. The high levels of fecal *Salmonella* Typhimurium observed at timepoints when the microbiota and SCFA levels were still depleted also indicate that microbiota-related factors play an important role in resistance against colonization by this pathogen.

### Flagellar motility is not required for *Salmonella* Typhimurium to colonize the gut or translocate to mesenteric lymph nodes after bowel prep in mice

Normally, flagellar propulsion is critical for *Salmonella* Typhimurium and other bacterial pathogens to penetrate the mucus layer of the gut and infect the underlying epithelium^33,34^. However, given the defects in the cecal mucus layer that we observed 6 hours after bowel prep treatment (**Fig. 3B**), we hypothesized that a non-motile *Salmonella* Typhimurium mutant might be able to colonize mice that had undergone bowel prep. To test this hypothesis, we challenged mice with wild-type *Salmonella* Typhimurium or a non-motile *Salmonella* Typhimurium Δ*flhD* mutant 6 hours after bowel prep or vehicle treatment. The Δ*flhD* mutant is missing the gene responsible for control of flagellar production^35^ but, similar to wild-type *Salmonella* Typhimurium, is resilient to osmotic perturbation (**Fig. S4**). Consistent with our hypothesis, mice treated with bowel prep and inoculated with the *Salmonella* Typhimurium Δ*flhD* mutant were efficiently colonized by the pathogen, whereas vehicle-treated mice were not (**Fig. 4A**).

**Figure 4.**
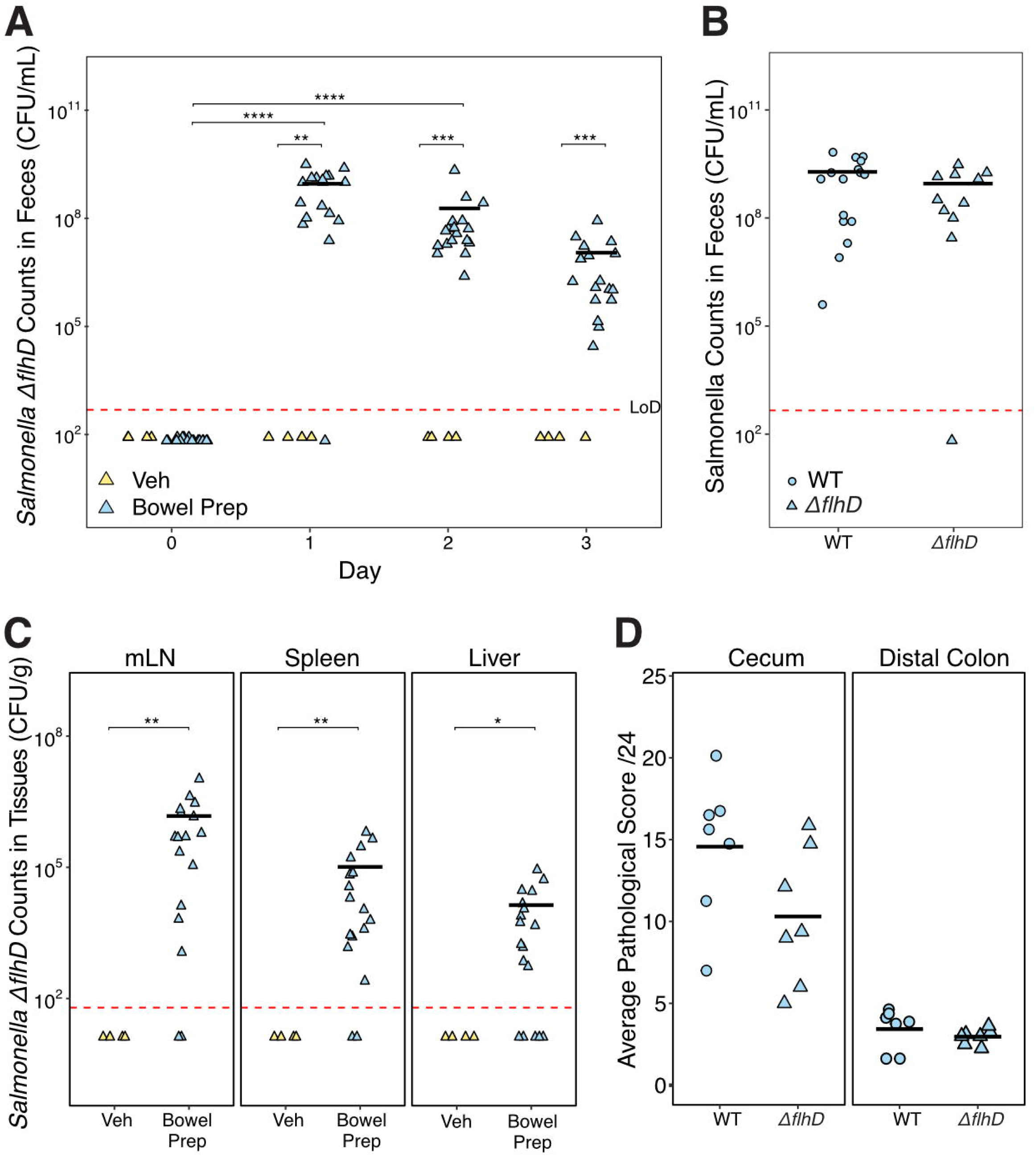
Flagellar motility is not required for *Salmonella* Typhimurium mutant (Δ*flhD*) gut colonization, translocation to the mesenteric lymph nodes, or pathology in mice subjected to bowel prep. (A) *Salmonella* Typhimurium Δ*flhD* colonization levels in mice inoculated 6 hours post bowel prep- or vehicle-treatment (Veh *n*=4, Bowel Prep *n*=17, two independent experiments for Bowel Prep, one for Veh). Pathogen counts were measured by plating feces over a 3-day period after inoculation. Differences in timepoints were analyzed Wilcoxon ranked-sum test, and differences between bowel prepped-samples at different timepoints were analyzed with a Friedman test followed by Nemenyi post-hoc test. (B) Wild-type (WT) and *Salmonella* Typhimurium Δ*flhD* counts in the feces one day after inoculation (WT *n*=16, Δ*flhD n*=17, two independent experiments). The differences between the treatment groups were measured using the Wilcoxon ranked-sum test. (C) *Salmonella* Typhimurium Δ*flhD* levels in mesenteric lymph nodes (mLN), spleen and liver (Veh *n*=4, Bowel Prep *n*=17, two independent experiments for Bowel Prep, one for Veh). Translocation in organ tissue was measured 72 hours after bowel prep. (D) Histopathological scores of the distal colon and cecum sections from bowel prepped mice (WT *n*=7, Δ*flhD n*=7). **Statistics:** Differences between treatment groups were measured using the Wilcoxon ranked-sum test. p > 0.05; ns (not significant, not shown), p < 0.05; *, p < 0.01; **, p < 0.001; ***, p < 0.0001; ****. **Abbreviation**: LoD, limit of detection.

In addition, among bowel prep-treated mice, fecal levels of the Δ*flhD* mutant were comparable to those of wild-type *Salmonella* Typhimurium one day after inoculation (**Fig. 4A,B**). Finally, the Δ*flhD* mutant was able to translocate from the gut to the mesenteric lymph nodes, liver, and spleen in bowel prep-treated mice but not vehicle-treated mice (**Fig. 4C**). These observations indicate that bowel prep–induced disruption allows *Salmonella* Typhimurium to colonize the gut and translocate to extra-intestinal organs without the normal requirement for motility. Consistent with this, H&E histopathology scores did not significantly differ between mice infected with the Δ*flhD* mutant and those infected with the wild type in the context of bowel prep (**Fig. 4D**), indicating that loss of motility did not significantly alter the overall pathology under these conditions, although a non-significant trend was observed. This supports the idea that the loss of the mucus barrier after bowel prep may reduce the necessity of flagellar motility.

### Bowel prep induces a small but coordinated cecal tissue transcriptional response

Given that both wild-type and non-motile *Salmonella* Typhimurium were able to colonize and disseminate following bowel prep, we next asked whether the host tissue responses might contribute to this increased susceptibility. We performed bulk RNA sequencing and analyzed the transcriptome of mouse cecal tissue at baseline and 6 hours post-bowel prep. We hypothesized that gene expression of host mucus and barrier functions genes might be affected after bowel prep, given the depletion of the mucus barrier (**Fig. 1C,D**). Groups were not significantly separated in principal component analysis (PCA) clustering of the cecal gene expression of bowel prep-treated mice (**Fig. 5A**) (R^2^ = 0.14, p = 0.098), or dispersed (p = 0.492) based on treatment. Likewise, expression of individual genes six hours post-bowel prep was largely unchanged from the baseline, with only 25 genes significantly differentially expressed (**Fig. 5B**). Of the top 50 differentially expressed genes by absolute Z score, few were related to mucus production and barrier function (**Fig. S5A**), and individual genes related to mucus production, barrier function, and immune system processes showed no significant changes between baseline and bowel prep (**Fig. S5B**).

**Figure 5.**
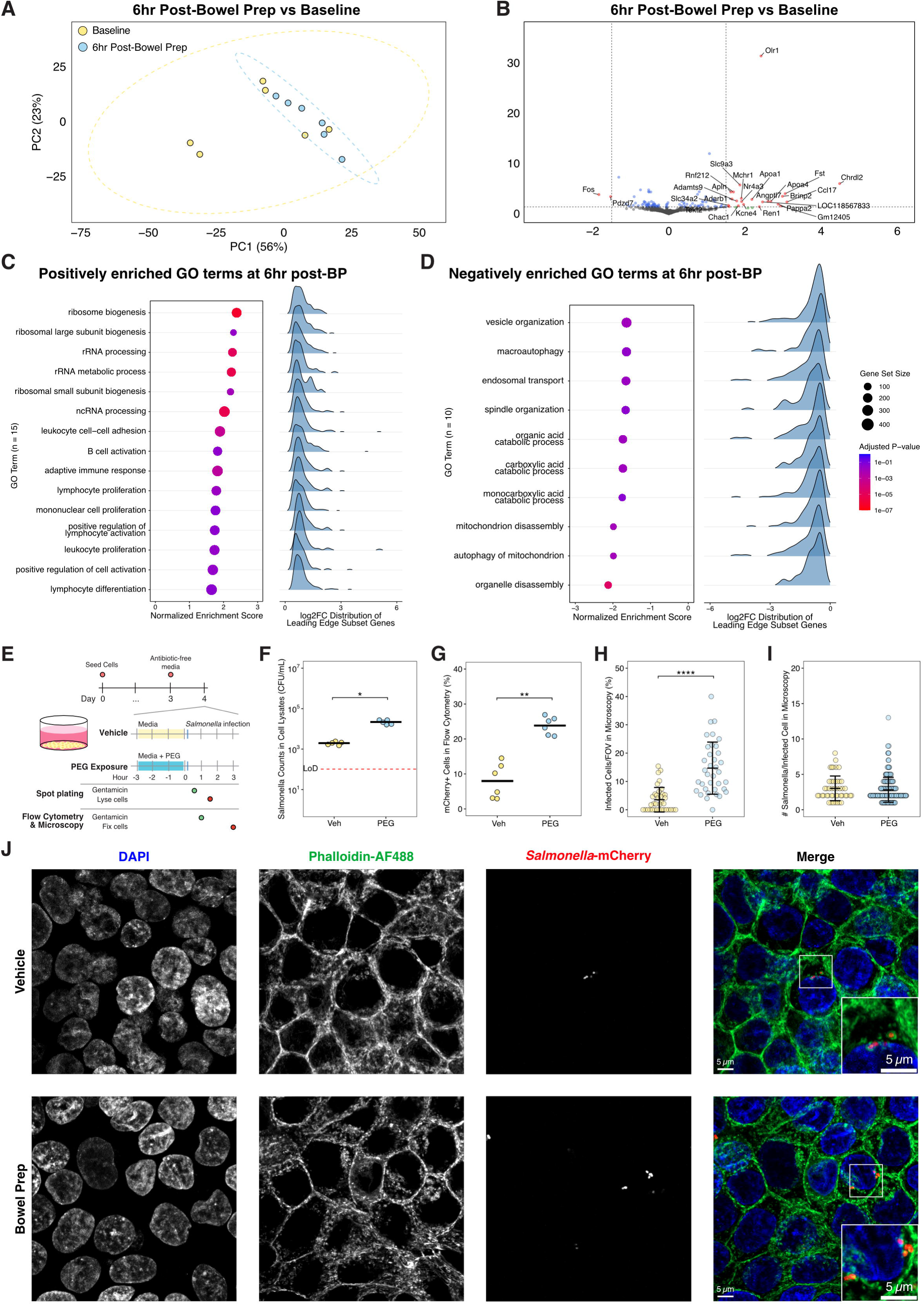
Bowel prep induces mild changes in epithelial gene expression *in vivo*, while PEG promotes *Salmonella* Typhimurium invasion *in vitro*. (A) Principal component analysis (PCA) plot of cecal gene expression data between mice at baseline and 6 hours post-bowel prep treatment. Group separation was assessed by PERMANOVA and group dispersion by PERMDISP. (B) Volcano plot of differential gene expression in mouse cecal tip tissue at 6 hours after bowel prep compared to baseline. The plot shows the names of genes with adjusted p-values (Benjamin-Hochberg false discovery rate–corrected) < 0.05 and Log2 fold change > 1.5. (C-D) Gene set enrichment analysis (GSEA) of Gene Ontology (GO) Biological Process terms on differentially expressed genes at 6-hr post-bowel prep vs baseline. Dot plots display normalized enrichment scores (NES), with point size representing gene set size and color indicating adjusted p-values (Benjamin-Hochberg false discovery rate–corrected), while ridge plots show the log_2_ fold change distributions of leading-edge subset genes in each gene set. (C) Significantly positively enriched GO terms (NES > 0). (D) Significantly negatively enriched GO terms (NES < 0). (E) Schematic of *in vitro* model of *Salmonella* Typhimurium invasion in HT-29 monolayers after 3 hours of PEG exposure. For spot plating, cells were treated with gentamicin for 1 hour after 30 minutes of initial infection with *Salmonella* Typhimurium, then lysed. For flow cytometry and microscopy, cells were treated with gentamicin for 2 hours after 1 hour of initial infection with mCherry-*Salmonella* Typhimurium, then fixed. (F) Spot plating *Salmonella* Typhimurium counts in cell lysates (*n*=5 for each condition). Differences in bacteria counts were analyzed with a Wilcoxon Ranked-Sum test. (G) Proportion of cells with mCherry-positive signal from flow cytometry (*n*=6 per condition from two independent experiments). Infected cell proportions were analyzed with a Wilcoxon Ranked-Sum test. (H) Count of infected cells per field of view (FOV) from microscopy images (*n*=32 FOVs per condition from 3 independent experiments). Differences in *Salmonella* Typhimurium infection across conditions were analyzed with a t-test, with black bars showing the mean and standard deviation. (I) *Salmonella* Typhimurium counts in infected cells (Veh *n*=46, Bowel Prep *n*=193 from the cells in [H]). t-test, black bars showing the mean and standard deviation. (J) Representative confocal micrographs of fluorescently labelled HT-29 cells infected with mCherry-*Salmonella* Typhimurium (*red*). Cells were fixed and stained with DAPI (*blue*) to visualize nuclei and Alexa Fluor 488–conjugated phalloidin (*green*) to label filamentous actin in vehicle- and PEG-treated conditions. Insets show magnified views of boxed regions highlighting *Salmonella*-infected cells. **Statistics**: p > 0.05; ns (not significant, not shown), p < 0.05; *, p < 0.01; **, p < 0.001; ***, p < 0.0001; ****. **Abbreviation**: LoD, limit of detection.

We then performed pathway-level analysis to capture coordinated biological responses through Gene Set Enrichment Analysis (GSEA). We discovered that at 6 hours post-bowel prep, 15 Gene Ontology (GO) Biological Process terms were significantly positively enriched (Normalized Enrichment Score, NES > 0), including terms related to protein synthesis and adaptive immune system functions (**Fig. 5C, Table S3**). On the other hand, we found 10 significantly negatively enriched GO terms (NES < 0), which included organelle recycling processes, intracellular trafficking processes, and catabolic functions (**Fig. 5D, Table S3**). Positively enriched GO terms shared overlapping core enrichment genes, largely among pathways related to protein synthesis and immune activation (**Fig. S5C**), while negatively enriched GO terms overlapped in genes associated with organelle degradation and catabolic processes (**Fig. S5D**). Although individual fold-changes were modest, the significant NES indicate a coordinated shift in gene expression across these pathways.

Together, these analyses suggest that bowel prep triggers an early but modestly scaled, coordinated transcriptional response in cecal tissue, characterized by upregulation of protein synthesis and immune pathways alongside downregulation of catabolic processes.

### PEG exposure promotes *Salmonella* Typhimurium invasion in a human intestinal epithelial cell model

Because the transcriptional response in cecal tissue suggested only modest host changes, we next asked whether bowel prep conditions could directly alter epithelial susceptibility to bacterial invasion using *in vitro* human intestinal epithelial cell models. To investigate this, we treated confluent monolayers of HT-29 cells, an established *in vitro* model of the human intestinal epithelium^36,37^, with PEG-supplemented media for 3, 6, or 24 hours (**Fig. S6A**). Given that cell culture media osmolality was 350 mOsm/kg, we adjusted *in vitro* osmolality to 600 mOsm/kg using PEG, mirroring the increase observed *in vivo* between vehicle and bowel prep-treated mice (**Fig. 1B**), in addition to testing higher osmolalities at 900 and 1200 mOsm/kg. We then infected monolayers in these conditions with *Salmonella* Typhimurium for 90 minutes, and counted intracellular bacteria from cell lysates. Invasion was maximal at 600 mOsm/kg condition, with intracellular counts increasing 6.9-fold from 1 hour (4 x 10^4^ CFU/mL) to 3 hours (2.75 x 10^5^ CFU/mL) of pre-treatment (**Fig. S6B**). Compared to the vehicle condition, monolayers treated at 600 mOsm/kg for 3 hours before infection showed an 11.6-fold increase in mean *Salmonella* Typhimurium counts (1.9 x 10^3^ vs 2.2 x 10^4^ CFU/mL, p = 0.01193) (**Fig. 5F**). These results indicate that short-term exposure to PEG substantially increases epithelial susceptibility to bacterial entry, even in the absence of the mucus layer and microbiota.

To further characterize the degree of intracellular invasion, we treated HT-29 monolayers for 3 hours at 600 mOsm/kg and infected cells with a constitutive mCherry-expressing *Salmonella* Typhimurium strain^38^ for another 3 hours to allow bacteria expansion (**Fig. 5E**). We hypothesized that since there was more *Salmonella* Typhimurium in cell lysates of PEG-treated cells, there would be increased mCherry-*Salmonella* Typhimurium at the single cell level. Indeed, flow cytometry showed a threefold increase in the proportion of infected cells relative to vehicle controls (23.81% vs 7.98%, p = 0.0022) (**Fig. 5G, Fig. S6C,D**). Confocal microscopy corroborated these findings: PEG–treated monolayers contained 4.2 times more infected cells than the vehicle controls (14.69% vs 3.54%, p = 2.78 x 10^-8^) (**Fig. 5H, Fig. S6E**). In contrast, the intracellular bacterial load per infected cell was similar across conditions (**Fig. 5I**), suggesting that bowel prep-conditions may enhance bacterial entry but not intracellular replication once invasion occurs. Finally, phalloidin staining revealed modest actin cytoskeleton disorganization after PEG treatment, a phenotype that warrants further investigation (**Fig. 5J**).

To complement our monolayer invasion assays, we also tested the effects of bowel prep conditions in a gut-on-a-chip model that more closely recapitulates intestinal physiology^39^. In this system, bowel prep treatment modestly increased epithelial permeability and *Salmonella* Typhimurium translocation compared to controls, although the results did not reach statistical significance (**Fig. S6F–I**). Together, our results suggest that exposure to PEG enhances epithelial vulnerability to *Salmonella* Typhimurium invasion and translocation *in vitro*, phenotypes which could be contributing to the increased pathogen colonization and translocation *in vivo*.

### Bowel prep exacerbates colitis and enhances pathobiont translocation in human IBD microbiota-associated mice

In previous work, we systematically profiled the tolerance of 92 representative human gut bacterial strains to a range of osmotic conditions^19^. We found that many abundant commensals, particularly strict anaerobes, exhibited markedly reduced growth at osmolalities comparable to those induced by bowel prep; conversely, Enterobacteriaceae members were among the most osmotolerant taxa^19^. Given that *Salmonella Typhimurium* is highly osmotolerant and can disseminate after bowel prep, we next asked whether IBD-associated pathobionts might similarly thrive under these conditions. We reasoned that Enterobacteriaceae-derived pathobionts present in the microbiota of patients with ulcerative colitis might be particularly well suited to exploit the altered gut environment following bowel prep. To test this, we examined the ability of 130 strains isolated from patients with UC that belonged to genera previously identified as pathobionts^40,41^ (family Enterobacteriaceae, genera: *Proteus, Morganella, Kluyvera, Klebsiella, Escherichia, Enterobacter* and *Citrobacter*) for their growth across a range of osmolalities simulating bowel prep. These potential pathobionts showed strong growth at osmotic levels experienced in the mouse gut post-bowel prep (∼765 mOsm/Kg) in both anaerobic and aerobic conditions (**Fig. S7A,B**). As expected, most strains were able to grow above 1100 mOsm/Kg, well above the levels many commensal bacteria are able to grow in^19^. These results indicate that pathobiont genera can thrive under the high osmotic conditions created by bowel prep.

Next, we asked whether the pathobionts’ relatively high osmotic resistance could lead to a shift in community composition under conditions of continued elevated osmotic stress. We cultured a fecal sample obtained from an individual with ulcerative colitis (hereafter referred to as hIBD1) under osmotic stress levels simulating normal (400 mOsm/Kg) and bowel prep-treated (800 mOsm/Kg) conditions. 16S rRNA sequence data showed a shift in community composition under the high osmolality condition (**Fig. S7C**), including increased abundance of the family Enterobacteriaceae. Specifically, the abundance of the genera *Morganella* and *Escherichia-Shigella* increased (**Fig. S7D**), indicating that even in a complex community, these potential IBD-associated pathobionts had a competitive advantage. The high osmotolerance of these UC-derived potential pathobionts, combined with the commensal depletion and barrier disruption observed post-bowel prep, suggests that they may have a competitive advantage *in vivo* under these conditions. We therefore next tested whether bowel prep could promote the expansion and systemic dissemination of these bacteria in a mouse model of ulcerative colitis.

We first humanized germ-free mice with fecal samples from a healthy participant (hHealthy) or the characterized patient with ulcerative colitis (hIBD1) and found that there was no observable long-term translocation 2 weeks post-bowel prep (**Fig. S8A,B**). This suggests that bowel prep by itself does not cause sustained bacterial translocation. Given reports of post-colonoscopy flare-ups in patients with IBD, we hypothesised that acute inflammation might affect the response to bowel prep in an IBD-microbiota context. To test this, we colonized mice with two patient ulcerative colitis microbiota and a healthy participant microbiota (hIBD1, hIBD2 and hHealthy, respectively), and induced acute colitis by treating mice with 2% dextran sodium sulphate (DSS) in drinking water for 5 days (**Fig. 6A, S8C**). We measured disease activity index (DAI, a composite score of body weight loss, stool consistency, fecal bleeding, and behaviour^42–44^) daily starting from the onset of DSS treatment. Both hIBD groups showed higher DAI than the healthy controls, with variability between donors, consistent with the microbiota-dependence of DSS response^45,46^ (**Fig. S8D,E)**.

**Figure 6.**
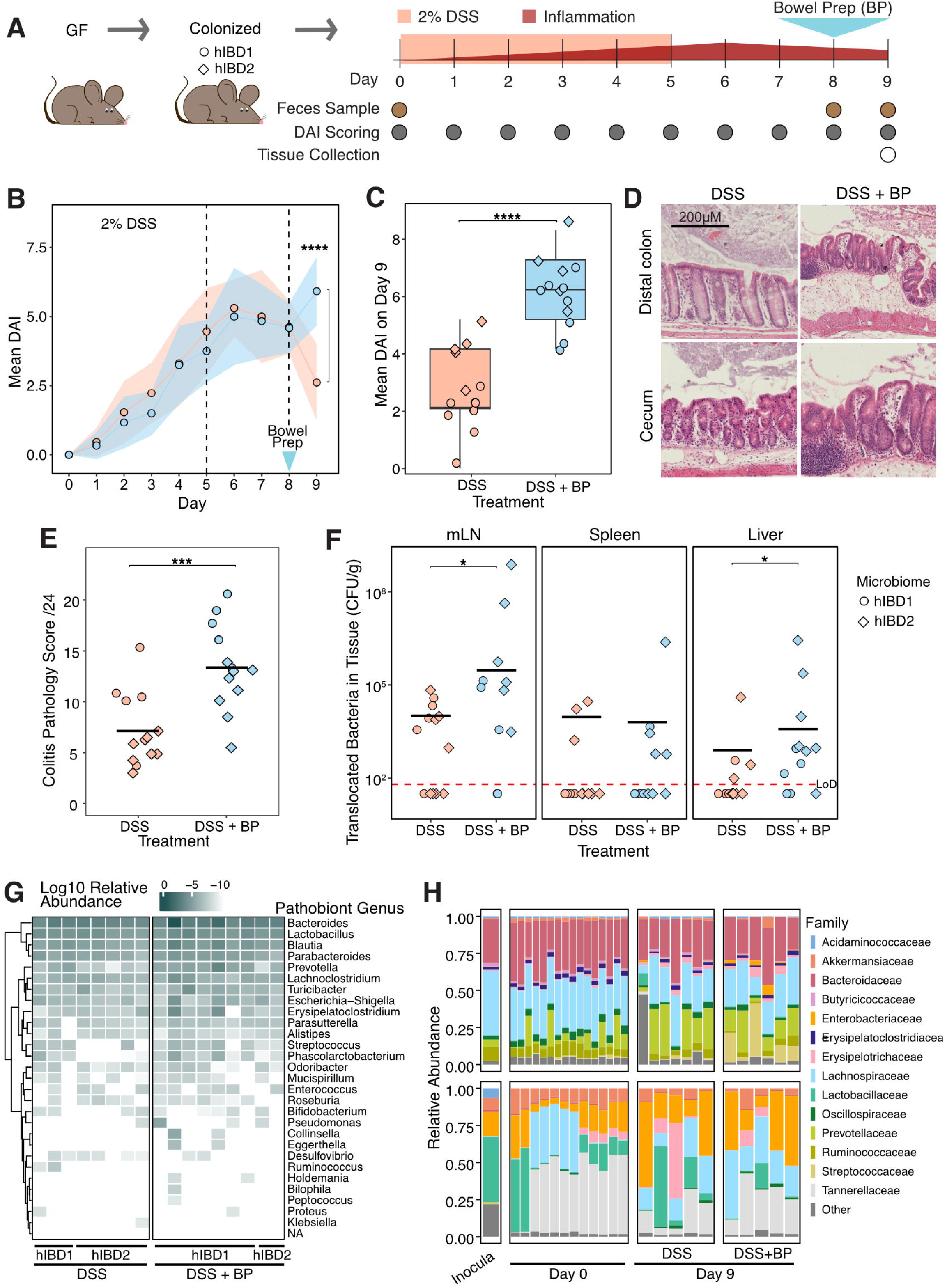
In human IBD microbiota-associated mice undergoing acute inflammation, bowel prep exacerbates colitis and increases extra-intestinal bacterial translocation. (A) Schematic of human microbiota colitis model. Germ-free mice were inoculated with a fecal sample from one of two human ulcerative colitis patients. After microbiota stabilisation, mice were treated with 2% DSS in drinking water for five days, received bowel prep on day 8, and were euthanized on day 9. Disease activity index (DAI) was assessed daily, and tissues were collected on day 9 for histology. (B) Mean colitis DAI was significantly different between no-treatment control and bowel-prep treated mice only on day 9, 24 hours after bowel prep (BP) (control *n =* 13, bowel prep *n* = 13). (C) Mean DAI scores of individual mice on day 9, from (B). (D) Representative H&E stained sections of the distal colon and cecum show increased immune infiltration, severity, and crypt damage in DSS-treated mice that received bowel prep, compared to ones that did not. (E) Colitis pathology score in the distal colon shows significantly worsened DSS-induced colitis in mice that received bowel prep. (F) Significant levels of bacterial translocation from the gut to the mesenteric lymph nodes (mLN) and liver, but not the spleen, were observed one day after hIBD mice received bowel prep (control *n =* 13, bowel prep *n* = 13). (G) 16S rRNA sequencing of bacteria translocated to the mesenteric lymph nodes of mice from (A) show the presence of genera from pathobionts associated with IBD^6,48–62^. (H) Stool 16S rRNA sequencing of mice from (A) colonized with hIBD1 (*top*) or hIBD2 (*bottom*) microbiotas before and after DSS treatment with and without bowel prep. LoD, limit of detection. p > 0.05; ns (not significant, not shown), p < 0.05; *, p < 0.01; **, p < 0.001; ***, p < 0.0001; ****.

Because mice colonized with a healthy microbiota are not expected to develop spontaneous intestinal inflammation, bowel prep was performed only in hIBD-colonized mice. Bowel prep was performed two days after cessation of DSS (**Fig. 6A**) to avoid treatment interference, and to better mimic the clinical scenario in which colonoscopy is performed shortly after inflammation. We found that hIBD mice treated with bowel prep showed a 2-fold increase in DAI 24 hours later compared to the no-treatment controls (p = 0.00013, **Fig 6B,C**). This effect was preserved between donor groups though the magnitude varied (**Fig. S8F**). We then measured tissue pathology by performing H&E staining and scoring^47^ in the cecal tip and distal colon. Supporting the DAI results, we found that mice that had received bowel prep had an average cecal pathology score of 13.3 compared to untreated mice which scored 7.1 (out of 24, p = 0.00041, **Fig. 6D,E**).

Having measured significant increases in inflammation due to bowel prep, we then tested translocation of bacterial species to extra-intestinal organs. We found that the amount of anaerobic extra-intestinal bacteria increased 30-fold in the mesenteric lymph nodes (p = 0.037) and 5-fold in the liver (p = 0.015) 24 hours post-bowel prep compared to mice that received no treatment (**Fig. 6F**). We then performed 16S rRNA sequencing of the mesenteric lymph nodes to identify which bacteria translocated. This analysis revealed many pathobiont species associated with IBD independent of bowel prep (**Fig. 6G, S8H**). While potential pathobiont species were found in both bowel prep and no treatment controls likely due to the DSS treatment, the number of observed species trended higher in the bowel prep group (p = 0.11, **Fig. S8G**). Finally, 16S rRNA sequencing of stool samples collected at the start and end of the experiment showed little change in microbiota composition, which remained largely reflective of their starting inocula (**Fig 6H**).

Together, these experiments demonstrate that bowel prep exacerbates colitis in human IBD microbiota-associated mice and enables osmotolerant Enterobacteriaceae pathobionts to expand and translocate beyond the gut, highlighting a mechanism by which transient environmental disruption can precipitate both local and systemic disease.

## Discussion

In this study, we set out to use mouse and *in vitro* models of bowel prep to determine whether the procedure increases the gut’s vulnerability to colonization by pathogens. We showed that bowel prep with PEG (1) alters the intestinal environment (**Fig. 1)**, (2) eliminates the natural microbiota’s and epithelial protection against *Salmonella* Typhimurium invasion (**Figs. 2–5**), and (3) exacerbates colitis in a human IBD microbiota-associated mouse model (**Fig. 6**). Our findings show that bowel prep reduces the gut microbiota’s natural resistance to pathogens through multiple mechanisms, including increased levels of osmotic stress, depletion of the mucus layer, and reduced competition by commensal bacteria.

Shortly after bowel prep is performed, before the commensal microbiota and gut environment have a chance to recover, hosts are extremely vulnerable to pathogens. In this study, a inoculation dose as low as 1,000 *Salmonella* Typhimurium cells successfully colonized the gut of mice treated with bowel prep 6 hours earlier and disseminated to the lymph nodes, spleen, and liver (**Fig. 2G,H**). This dose is between ten thousand and a million times lower the usual dose used in the streptomycin models of *Salmonella* Typhimurium infection^63^. This low-dose infection occurred despite the absence of bicarbonate treatment, which is commonly used to ensure the *Salmonella* Typhimurium load is not negatively affected by stomach acidity^64^, or pretreatment with antibiotics^23,24^. While these findings stem from controlled preclinical models, they reveal that acute disruption of the gut environment alone is sufficient to reduce colonization resistance against enteric pathogens.

We observed the highest levels of *Salmonella* Typhimurium colonization when inoculation occurred 6 hours post-bowel prep, when the mucus layer was at its thinnest (**Fig. 3B,F**). These findings align with previous work in mice and humans with ulcerative colitis showing that thinning of the intestinal mucus layer enhances bacterial penetration of the gut epithelium and increases host susceptibility to pathogen infection^30,65^. Consistent with this, mice genetically engineered to lack intestinal mucus (*Muc2* -/-) display increased susceptibility to *Citrobacter rodentium*^66^ as well as *Salmonella* Typhimurium^67^.

Normally, *Salmonella* Typhimurium needs a functioning flagellum to colonize the mammalian gastrointestinal tract^64^, and previous studies have reported that disruption of *Salmonella* Typhimurium motility reduces pathogen fitness and invasiveness *in vitro* and *in vivo*^68–70^. However, in absence of mucus post-bowel prep, the non-motile *Salmonella* Typhimurium Δ*flhD* mutant was able to colonize and translocate to the mesenteric lymph nodes (**Fig. 4C**). As the FlhD protein, a master regulator of flagellar biogenesis and motility, is involved in coordinating virulence^71,72^, the ability of *Salmonella* Typhimurium Δ*flhD* to translocate suggests other virulence factors may facilitate *Salmonella* Typhimurium translocation after bowel prep. Specifically, type III secretion systems (T3SS) encoded by *Salmonella* pathogenicity islands 1 and 2 (SPI-1 and SPI-2) are independent of FlhD regulation, and enable *Salmonella* Typhimurium to colonize, invade, and survive in the host without requiring motility^73,74^. This finding expands our understanding of pathogen-host interactions in disrupted gut environments and suggests that non-motile pathogens may also be able to exploit these conditions to invade the gut mucosa.

Although mucus loss occurred throughout the mouse digestive tract, we observed pathology in *Salmonella* Typhimurium infected cecal tissues but not in the colon (**Fig. 2E,F**), unlike streptomycin-treated mice (**Fig. S2E,F**). While a previous study suggested the cecum is the preferred side of *Salmonella* Typhimurium infection due to incomplete mucus coverage^33^, our findings suggest that the mechanical changes induced by bowel prep, particularly the thinning of the cecal mucosa, may further enhance tissue invasion at this location. Supporting this, a recent study has found that hypertonic stimulation increases bacterial adhesion to host cells *in vitro*^75^.

Beyond the effects of the mucus, our data from mice suggest that microbiota depletion is a key driver of post-bowel prep susceptibility to pathogens. After 24 hours, osmolality and mucus had recovered, but luminal microbiota richness and SCFA levels remained depleted (**Fig. 3A–D**), and *Salmonella* Typhimurium was still able to colonize the gut (**Fig. 3E–G**). Higher SCFA levels, particularly butyrate, have been linked to lower *Salmonella enteritidis* burden in chickens through induction of host defense peptide gene expression^76^, while depletion of butyrate-producing Clostridia increases aerobic expansion of *Salmonella* Typhimurium in mice^77^. Given the observed reduction in intestinal butyrate levels following bowel prep, it is possible that *Salmonella* Typhimurium experiences less inhibition of its virulence, which could contribute to its ability to invade tissue and translocate to extraintestinal organs (**Fig. 3E–G**). This aligns with a previous mechanistic study showing that microbiota-derived butyrate acylates and inactivates HilA, the master transcriptional activator of SPI-1, leading to less colonization in mice compared to an acylation-deficient mutant^78^.

Interestingly, our *in vitro* data suggests that PEG exposure alone can increase epithelial susceptibility to invasion, but not intracellular expansion (**Fig. 5E–J**). This finding is consistent with prior reports showing that PEG-adjusted media decreases HT-29 cell viability and growth within a day of culturing^79^, while higher osmolality media (mannitol-supplemented) disrupts tight junctions of Caco-2 intestinal epithelial cells^80^. Consistent with this interpretation, we observed thinning of the cecal mucosa 6 hours after bowel prep *in vivo* (**Fig. 1E**), suggesting that the treatment may directly damage epithelial cells and weaken the barrier. On the bacterial side, osmotic stress has been shown to increase invasion and adherence phenotypes across multiple *Salmonella* strains into intestinal epithelial cell lines^81–83^, as well as upregulation of virulence factors such as Sip proteins and flagellin^84^. Similarly, in other enteric pathogens like *Shigella* and enterohemorrhagic *E. coli* O157:H7, high-osmolality conditions trigger increased expression of T3SS structural components and effector proteins ^85,86^. Thus, the combination of host epithelial disruption and pathogen hyperinvasiveness under osmotic stress–as observed during bowel prep– likely act synergistically, amplifying pathogen exploitation of host vulnerability in the absence of normal colonization resistance.

Supporting the rapid changes in pathogen invasion *in vitro*, our mouse data further indicate that bowel prep increases susceptibility to infection within hours, yet the host mounts only a limited response during this acute window (**Fig. 5A,B**). At 6 hours post-bowel prep, when pathogens susceptibility peaked, cecal mucosa transcriptomes showed only minor changes. This suggests that epithelial cells are poorly equipped to counteract this perturbation within the timescale of maximal vulnerability. By contrast, other large-scale gut perturbations evoke stronger and faster transcriptional response. For instance, *Salmonella* Typhimurium infection upregulates cytokine gene expression within 2.5 hours in human intestinal organoids^87,88^, and genes related to the inflammatory response within 8 hours in the mouse colon^89^. Interestingly, we found a coordinated enrichment of adaptive immune system GO terms in GSEA 6 hours after bowel prep (**Fig. 5C,D**), but this modest activation was insufficient to prevent *Salmonella* Typhimurium invasion and dissemination.

Taken together, these findings show that bowel prep creates a short-lived but critical window in which both the host and invading pathogens are shifted toward a state of heightened susceptibility. While epithelial defenses are weakened and pathogens upregulate invasion pathways, an additional concern is that pathobionts already present within the microbiota—rather than only externally introduced pathogens like *Salmonella* Typhimurium—may also exploit this disruption. Importantly, we found that, unlike commensal bacteria, which display limited resilience to increased osmolality^19^, IBD-associated pathobionts are highly tolerant to the osmotic perturbation induced by bowel prep, and can expand under these conditions *in vitro* (**Fig. S7**). However, in germ free mice colonized with either IBD or healthy human microbiota, we detected no significant bacterial translocation two weeks after bowel prep (**Fig. S8A,B)**. These hIBD-colonized mice also showed no signs of colitis, suggesting that bowel prep in the absence of ongoing inflammation is not sufficient to exacerbate disease.

In contrast, when bowel prep was performed in the context of DSS-induced acute inflammation, both colitis severity and pathology were significantly worsened in IBD microbiota-associated mice (**Fig. 6D–E**). These mice also exhibited significantly increased extra-intestinal bacterial translocation relative to non-bowel-prep controls (**Fig. 6F**), mirroring the phenotype observed in *Salmonella* Typhimurium-challenged healthy mice. Sequencing of bacteria recovered from the mesenteric lymph nodes identified several IBD-associated pathobiont genera, including *Bacteroides*, *Lactobacillus*, and *Prevotella* (**Fig. 6G, S8H**). In addition to higher live bacterial counts in extra-intestinal sites, a greater proportion of bowel-prepped mice exhibited bacterial translocation compared to controls; however, the number of distinct species translocated per mouse did not differ significantly between groups (**Fig. S8G**). This further implies that while severe DSS-induced tissue damage may permit many species to penetrate the epithelium^65,90^, bowel prep likely increases the likelihood of such translocation events occurring.

Notably, the IBD microbiota associated with milder colitis during DSS treatment produced higher histopathology scores and greater bacterial translocation following bowel prep, suggesting that the impact of bowel prep is shaped by the specific microbiota context. Together, these findings highlight the interplay between bowel prep, inflammation, and microbiota composition, and suggest that future clinical studies should investigate how these factors jointly influence the risk of symptom exacerbation following the procedure^11–16^.

Overall, our findings reveal two complementary mechanisms by which bowel prep transiently lowers colonization resistance: on the pathogen side, disruption of the mucus barrier enables even non-motile *Salmonella* mutants to invade and disseminate, while on the microbiota side, osmotolerant Enterobacteriaceae pathobionts expand under PEG-induced osmotic stress and can translocate in the setting of inflammation. Together, these results define a discrete but clinically relevant window of heightened vulnerability that is acute and reversible, emerging within hours of bowel prep and persisting for at least 24 hours, but not observed after longer recovery. This temporal dynamic may provide a mechanistic basis for the observed post-colonoscopy flares in patients with IBD and underscores the importance of considering both host barrier integrity and microbiota composition when assessing patient risk; however, further clinical studies are necessary to validate and translate these findings. Furthermore, these insights highlight potential avenues to mitigate the adverse effects of bowel prep, such as the co-administration of SCFAs or barrier-preserving agents, or the development of alternative formulations designed to minimize epithelial and microbial disruption.

This study has several important limitations. Primarily, differences in gut physiology and microbiota composition between humans and mice could limit the application of our findings from mice to humans. Specifically, considerations such as transit time, size and physiology of the digestive tract (particularly the cecum, where the greatest *Salmonella* pathology was observed) will have an impact on the windows of susceptibility to pathogens in humans compared to mice. However, given the similarity of our model to clinical bowel prep (PEG laxative, removal of intestinal contents) and the support of our findings in *in vitro* human cell line system, we believe our findings to be applicable. Future studies of bowel prep in humans should investigate the presence of specific pathobionts in biopsies. Importantly, previous findings already highlight the increased representation of potential pathobionts members in the Enterobacteriaceae family such as *E. coli* and *Klebsiella* in IBD biopsies^91–93^. However, the presence of these genera, alongside a characterization of their potential ability to act as a pathobiont (for example via the presence of toxin genes) need to be correlated to health outcomes of the patients beyond the diagnosis at the time of colonoscopy. Still, particularly because of the technical challenges of measuring bacterial translocation in humans^11–16^, this animal model may provide important insights into which bacteria should be monitored in patients requiring more frequent colonoscopies, including those with IBD.

Similarly, differences between our *in vitro* model and the actual human gut may limit application of findings from this study system. However, in the bowel prep system, due to the much-reduced microbiota and mucus layer following the laxative treatment, this model is particularly apt to model pathogen infection in the context of PEG exposure. For instance, HT-29 cultures are an established model for studying pathogen interactions with the intestinal epithelium^36,37^. Although they contain a small subset of mucus-secreting cells^94^, they produce little mucus overall, reflecting the transient mucus-depleted state of the epithelium observed *in vivo* after bowel prep. Taken together, while the study systems used here have inherent limitations, their ability to mimic key aspects of bowel prep and pathogen susceptibility supports their utility in exploring the mechanisms underlying microbial translocation and infection.

In conclusion, this study establishes that, in both mouse and *in vitro* models, bowel prep rapidly alters the gut environment, increasing susceptibility to *Salmonella* Typhimurium and exacerbating inflammation in a colitis model. In addition, gut preparation promotes the translocation of intestinal bacteria from the gut to extraintestinal organs, such as nearby lymph nodes, the liver, and spleen. These findings suggest that gut preparation may have clinical implications that are not currently appreciated. Moreover, our findings suggest that other conditions that cause bouts of osmotic diarrhea, such as malabsorption and acute inflammation, could promote the growth and translocation of pathobionts that could further exacerbate disease states. While these findings provide important mechanistic insights, it is important to note that our results do not indicate a general risk to individuals undergoing clinically indicated bowel prep, but rather highlight potential vulnerabilities that should be investigated in specific at-risk populations such as IBD patients.

## Resource Availability

### Lead Contact

Carolina Tropini

## Acknowledgments

The authors acknowledge that the land we performed this research on is the traditional, ancestral, and unceded territory of the xwməθkwəy əm (Musqueam) Nation. The land our laboratory is situated on has always been a place of learning for the Musqueam people, who for millennia have passed on in their culture, history, and traditions from one generation to the next on this site. We encourage others to learn more about the native lands in which they live and work at https://native-land.ca/.

The authors thank Denise Monack, Aaron Dhanda, Christopher Lee, and members of the Tropini Lab for useful conversations and insight, in particular Michael Hunter for critically reading the manuscript and Angele Arrieta for ordering supplies and managing lab processes. We acknowledge the University of British Columbia Centre for Disease Modeling and Natalia Carranza Garcia for support with animal work. We thank Ben Swenor and William Bralower from Emulate for assistance with gut-on-a-chip experiments, Kunho Choi and the Vallance Lab for providing reagents and the mCherry-*Salmonella* Typhimurium strain, and Wanyin Deng and the Finlay Lab for providing *Salmonella* Typhimurium wildtype and mutant strains. This study also received support from the Life Sciences Institute Biofactorial High-Throughput Biology Core, supported by the University of British Columbia Global Research Excellence Biological Resilience Initiative. We thank Harper Health & Science Communications, LLC for editorial support of the initial manuscript submission. The authors acknowledge support from the Canadian Institute for Advanced Research / Humans and the Microbiome (FL-001253 Appt 3362, to C.T.), Michael Smith Foundation for Health Research Scholar Award (18239, to C.T.), Canada Foundation for Innovation / Infrastructure Operating Fund (38277), Canada Tier 2 Research Chair, Quantitative Microbiota Biology for Health Applications (CRC-2022-00036, to C.T.), Canadian Institutes of Health Research (CIHR) Project grant (PJT 191743), Crohn’s and Colitis Canada / Grants In Aid of Research (625155), Paul Allen Foundation / Allen Distinguished Investigators (ADI) Program (12935, to C.T. and S.R.S.), and the CH.I.L.D Foundation Chair in Pediatric Gastroenterology (to B.A.V.).

## Supplementary Figures

**Supplemental Figure 1.**
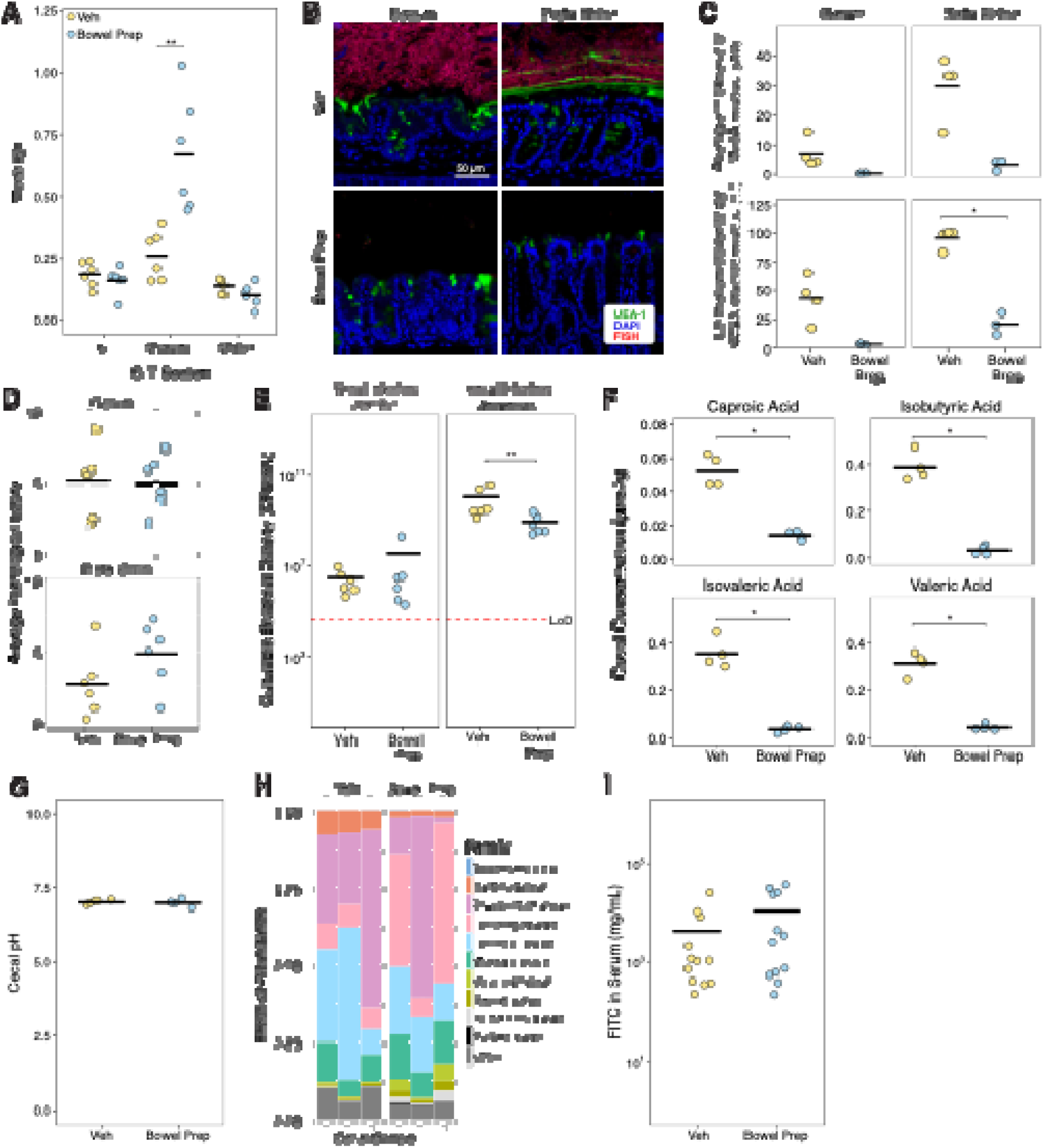
Bowel prep does not lead to overt pathology in the mouse gut at 6 hours post procedure, despite changes in cecal mass, mucus layer thickness, and short-chain fatty acid (SCFA) levels. (A) The mass of the cecum, the small intestine and colon measured in bowel prep- vs vehicle-treated mice (Veh *n*=6, Bowel Prep *n*=6). (B) Representative confocal micrographs of the cecal tip and distal colon (stained by FISH [*red*]) as well as UEA-1 stained mucus [*green*] following bowel prep. (C) Average thickness of mucus (*top*) and percentage of the epithelium covered (*bottom*), quantified using WGA fluorescence in bowel prep- vs vehicle-treated mice (Veh *n*=4, Bowel Prep *n*=3). (D) Average pathological scores of the cecum and distal colon in bowel prep- and vehicle-treated mice (Veh *n*=6, Bowel Prep *n*=6). (E) Bacterial loads in the small intestine (SI), as measured by culture under aerobic and anaerobic conditions (Veh *n*=6, Bowel Prep *n*=6). (F) Cecal SCFAs abundance in bowel prep- and vehicle-treated mice (Veh *n*=4, Bowel Prep *n*=4). (G) Cecal pH in bowel prep- and vehicle-treated mice (Veh *n*=4, Bowel Prep *n*=4). (H) 6 hours post-treatment, 16S rRNA sequencing of the contents of the cecum (Veh *n*=3, Bowel Prep *n*=3). These data are independent of those in Fig. 1G. (I) Gut permeability measured by FITC-Dextran movement from the gut into serum (Veh *n*=12, Bowel Prep *n*=12, two independent replicates). **Statistics**: Differences between treatment groups were analyzed using a Wilcoxon ranked-sum test. p > 0.05; ns (not significant, not shown), p < 0.05; *, p < 0.01; **, p < 0.001; ***, p < 0.0001; ****.

**Supplemental Figure 2.**
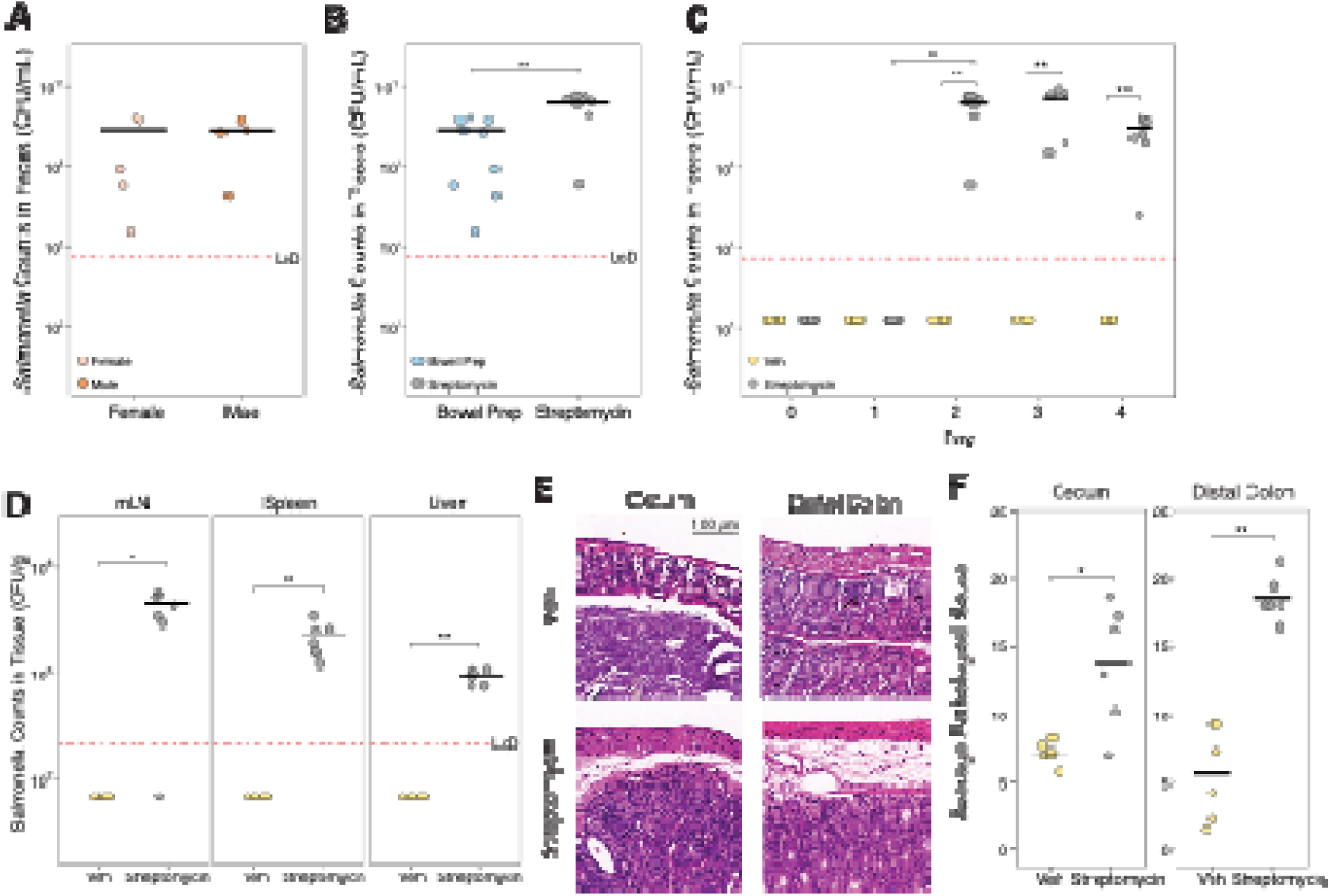
Comparison of two mouse models of *Salmonella* Typhimurium—pre-treatment with bowel prep or streptomycin—shows high levels of *Salmonella* Typhimurium burdens and robust translocation. (A) Fecal *Salmonella* Typhimurium levels 3-days post inoculation (Female *n*=5, Male *n*=4). Differences between treatment groups were analyzed using a Wilcoxon ranked-sum test. (B) Fecal *Salmonella* Typhimurium levels inoculated 6 hours post bowel prep (from **Fig. 2C**) and in streptomycin-treated mice 24-hours post inoculation (Bowel Prep *n*=9, Streptomycin *n*=7). Differences between treatment groups were analyzed using a Wilcoxon ranked-sum test. (C) Fecal *Salmonella* Typhimurium levels in vehicle- vs streptomycin-treated mice up to four days after inoculation (Veh *n*=5, Streptomycin *n*=7). Differences between timepoints were analyzed with a Wilcoxon ranked-sum test, and differences between Streptomycin samples were analyzed with a Friedman test followed by Nemenyi post-hoc test. (D) *Salmonella* Typhimurium translocation from the gut to the mesenteric lymph nodes (mLN), liver, and spleen 72 hours after inoculation (Veh *n*=5, Streptomycin *n*=7). Differences between treatment groups were analyzed using a Wilcoxon ranked-sum test. (E) H&E-stained sections of the cecum and distal colon in streptomycin- and vehicle-treated mice three days after inoculation with *Salmonella* Typhimurium. (F) Histopathological scoring of the distal colon and cecum in streptomycin- and vehicle-treated mice in cecum and distal colon at three days after inoculation with *Salmonella* Typhimurium (Veh *n*=6, Streptomycin *n*=6). Differences between treatment groups were analyzed using a Wilcoxon ranked-sum test. LoD, limit of detection. p > 0.05; ns (not significant, not shown), p < 0.05; *, p < 0.01; **, p < 0.001; ***, p < 0.0001; ****.

**Supplemental Figure 3.**
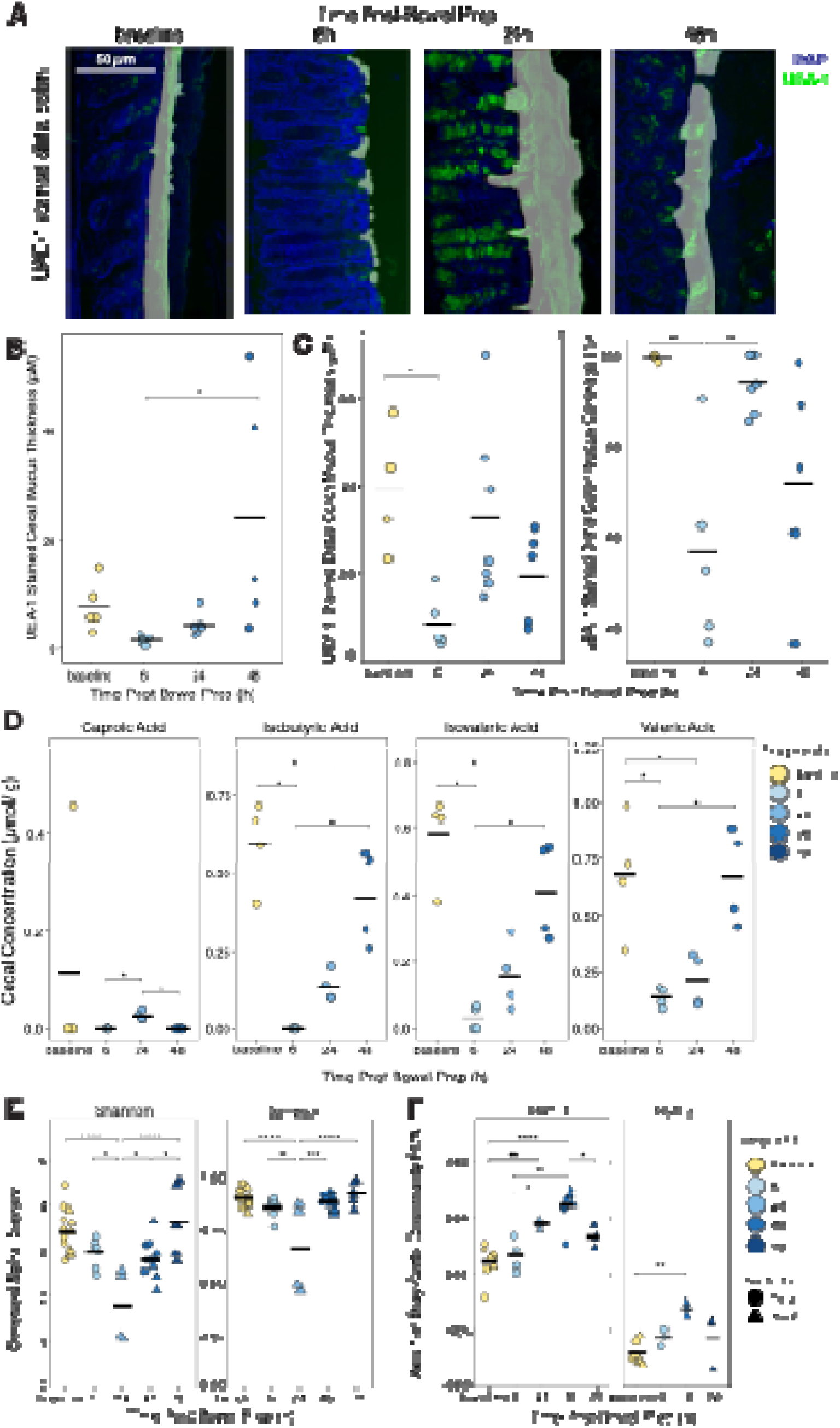
In the mouse cecum, short-chain fatty acid (SCFA) levels take longer to recover than the mucus layer and microbiota diversity after bowel prep. (A) Representative confocal images of the distal colon stained with DAPI which indicates host DNA (*blue*) and UEA-1 which indicates mucus (*green*). Contour lines generated in BacSpace software show the measured boundary used to calculate mucus thickness and coverage. (B) Mucus thickness in the cecum (baseline, 6 h, 48 h *n* = 5, 24 h *n* =6, 3 independent experiments). Differences between timepoints were analyzed using a one-way ANOVA with Tukey’s post-hoc test for multiple comparisons. (C) Mucus thickness and coverage in the distal 6 hours after bowel prep. Differences between timepoints were analyzed using a one-way ANOVA with Tukey’s post-hoc test for multiple comparisons. (D) SCFA levels at different timepoints after PEG treatment (each timepoint *n*=4). Differences between timepoints were analyzed with Kruskal-Wallis followed by Dunn’s post-hoc test. (E) Additional alpha diversity metrics in the cecal and fecal microbiome, determined from 16S rRNA sequencing (baseline *n*=14, 6h *n*=7, 24h *n*=4, 48h *n*=11, 72h *n*=6, 4 independent experiments). Differences between timepoints were analyzed using a one-way ANOVA with Tukey’s post-hoc test for multiple comparisons. (F) Bray Curtis dissimilarity values for the cecal and fecal microbiome, determined from 16S rRNA sequencing. Two Illumina sequencing runs are shown, with separate biological samples (Run 1 baseline *n*=8, 6h *n*=4, 24h *n*=4, 48h *n*=8, 72h *n*=3; Run 2 baseline *n*=6, 6h *n*=3, 48h *n*=3, 72h *n*=3). Differences between timepoints were analyzed using a one-way ANOVA with Tukey’s post-hoc test for multiple comparisons. p > 0.05; ns (not significant, not shown), p < 0.05; *, p < 0.01; **, p < 0.001; ***, p < 0.0001; ****.

**Supplemental Figure 4.**
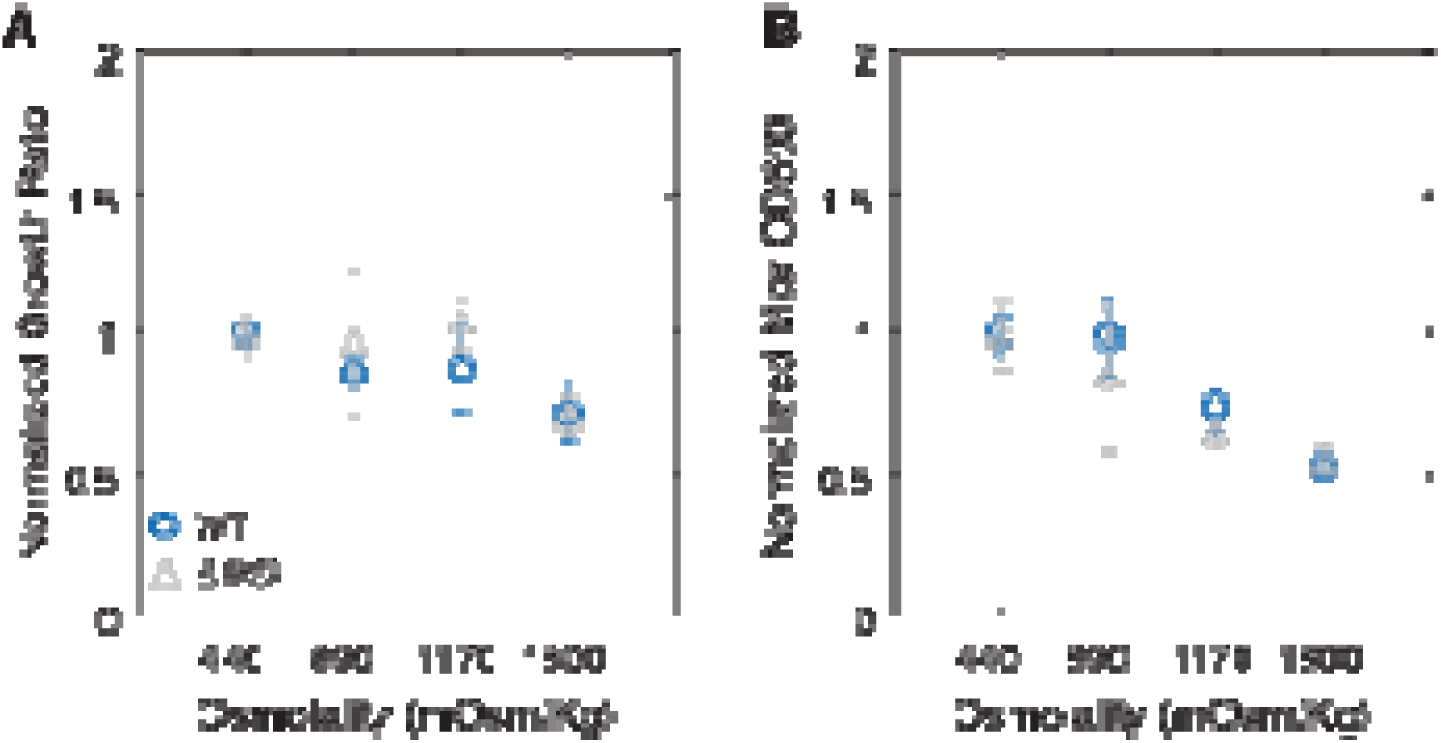
In culture, growth of the *Salmonella* Typhimurium non-motile Δ*flhD* mutant is robust under high osmolality conditions. (A) Normalized maximum growth rate and (B) normalized maximum optical density (OD600) of wild type and Δ*flhD Salmonella* Typhimurium in aerobic conditions and media of varying osmolalities, adjusted with PEG (each osmolality & strain *n*=4). Differences between timepoints and strains were analyzed using a two-way ANOVA with Tukey’s post-hoc test for multiple comparisons. p > 0.05; n.s. (not significant, not shown), p < 0.05; *, p < 0.01; **, p < 0.001; ***, p < 0.0001; ****.

**Supplemental Figure 5.**
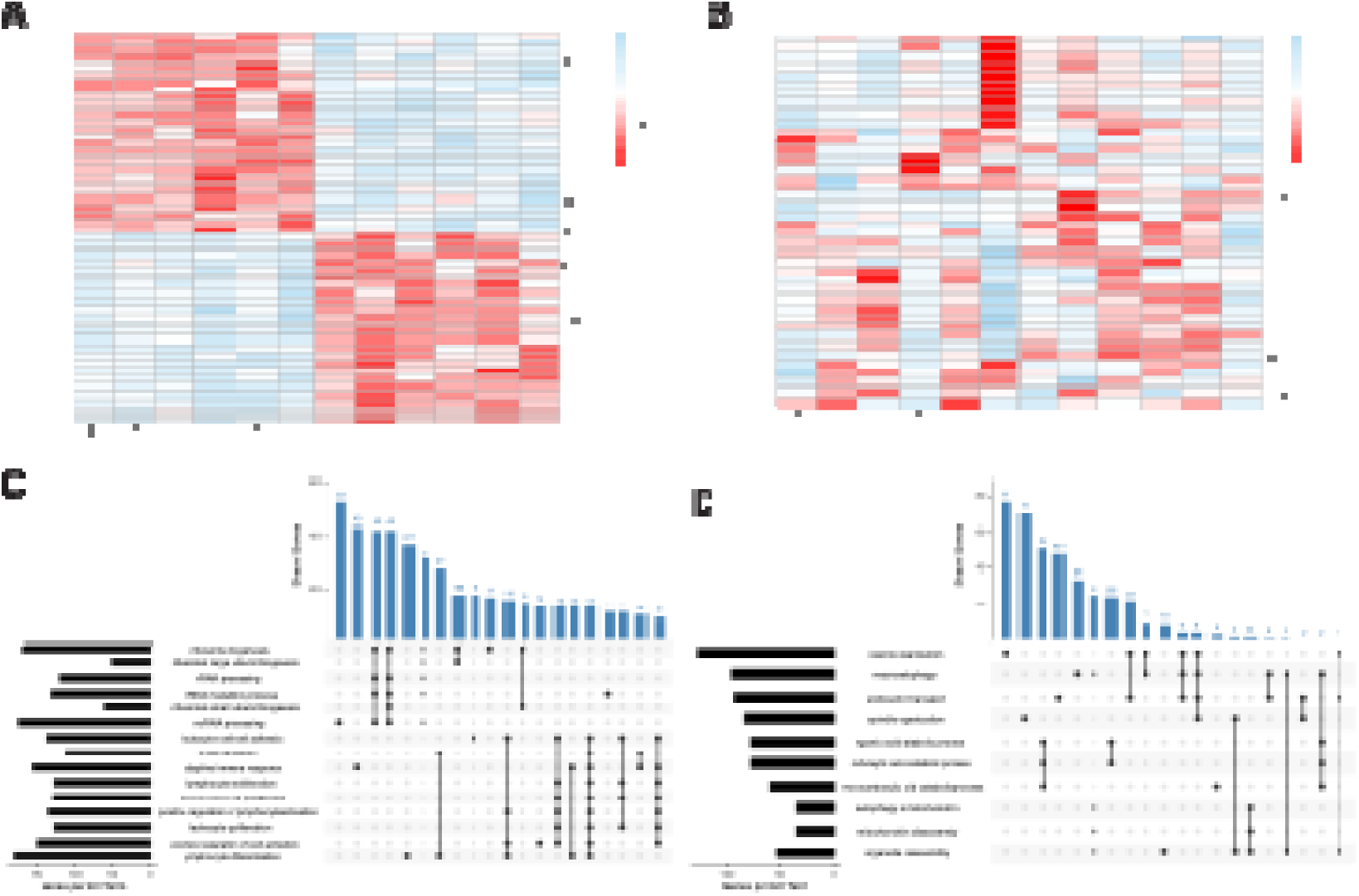
Bowel prep in mice does not lead to clear clustering of individual gene expression profiles, but enriched Gene Ontology (GO) terms share core enrichment genes. (A) Heatmap of the top 50 genes ranked by absolute Z score across all samples from cecal tip tissue at baseline and 6 hours after bowel prep. (B) Heatmap of 40 selected genes of interest, relating to mucus production, immune system activation, and tight junction formation. (C–D) Upset plots showing the overlap of leading-edge subset genes among significantly enriched Gene Ontology (GO) Biological Process terms identified by GSEA at 6hr post–bowel prep vs baseline (Table S3). The bar plot on the left indicates the number of core enrichment genes of each GO term; the bar plot above indicates the number of shared core enrichment genes between the combinations of GO terms connected in the matrix below. Only the top 20 shared gene overlaps are shown. (C) Significantly positively enriched GO terms (NES > 0) show shared gene membership among pathways related to protein synthesis and immune activation. (D) Significantly negatively enriched GO terms (NES < 0) show overlapping genes among pathways involved in organelle degradation and catabolic processes.

**Supplemental Figure 6.**
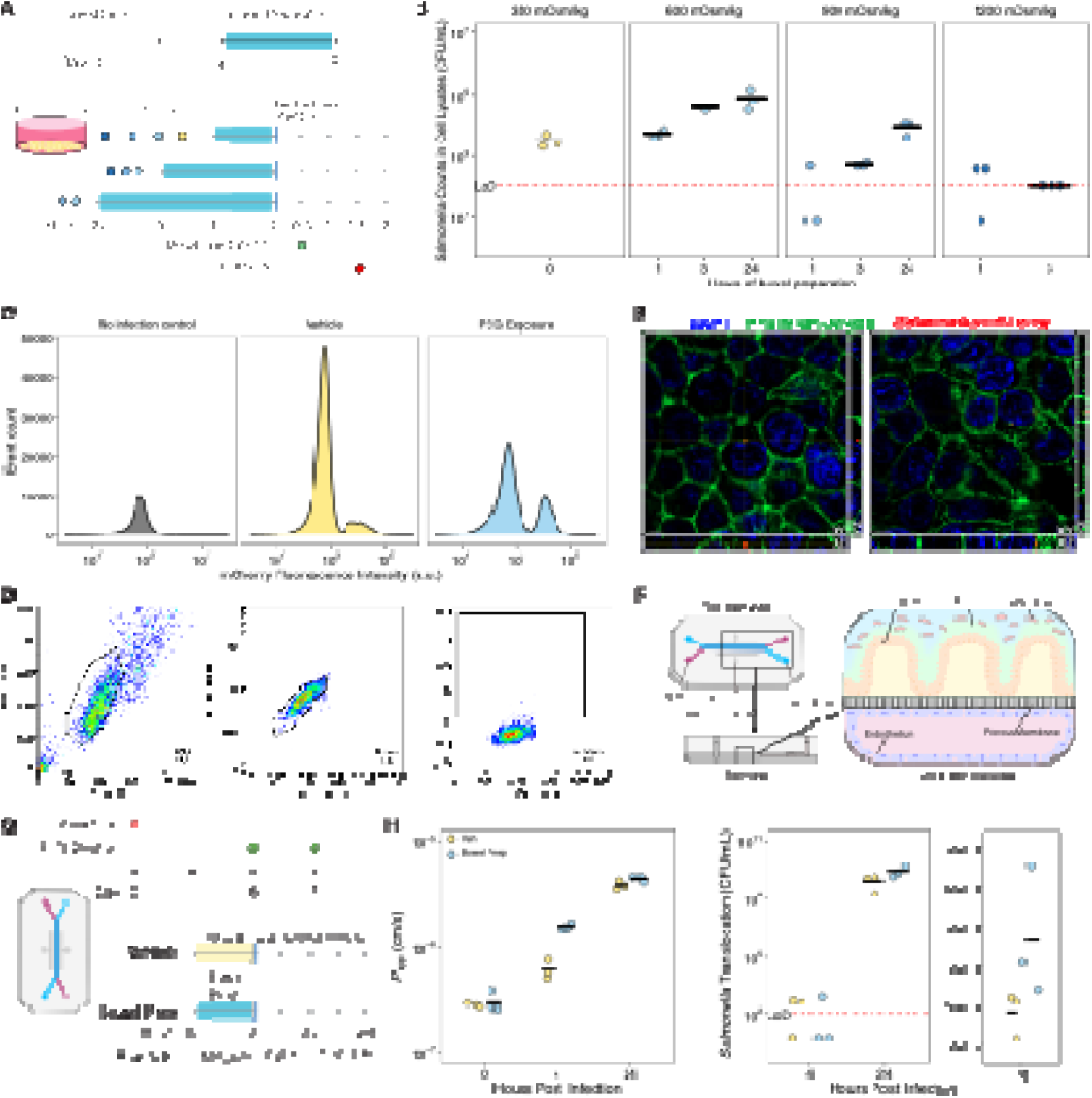
PEG exposure promotes *Salmonella* Typhimurium invasion in intestinal epithelial monolayers and increased permeability in a gut-on-a-chip model. (A) Schematic of bowel prep dosage experiment in HT-29 monolayers. Cells were exposed to media adjusted to 600, 900, and 1200 mOsm/kg using PEG for 1, 3, or 24 hours, then infected with *Salmonella* Typhimurium. After 30 minutes of infection, cells were treated with 100 µg/mL gentamicin for 1 hour, then lysed for spot plating. (B) HT-29 monolayers were treated at 600 or 900 mOsm/kg and infected with *Salmonella* Typhimurium; invasion was quantified by spot plating serial dilutions of cell lysates (n = 3 per condition per timepoint). Each dot represents one monolayer, with mean CFU/mL shown as black bars. (C) Representative flow cytometry histograms showing the mCherry fluorescence intensity event counts of the uninfected control (left, grey), infected vehicle–treated cells (middle, yellow), and infected bowel prep–treated cells (right, blue), from the experiment done in Fig. 5E. (D) Flow cytometry gating strategy to identify cells infected with mCherry-expressing *Salmonella* Typhimurium. The threshold for mCherry-positive signal was set to 1%> of events of the uninfected control population, as shown. (E) Orthogonal projections of HT-29 cells from Fig. 5J confocal micrographs stained with DAPI (blue) to visualize nuclei and Alexa Fluor 488– conjugated phalloidin (green) to label filamentous actin show that mCherry-*Salmonella* Typhimurium (red) is localized intracellularly. The XZ orthogonal view is shown at the bottom, and the YZ orthogonal view is shown on the right. The scale bars are 5 μm. (F) Schematic of gut-on-a-chip experiment designed to simulate the human gut environment, with top down, side, and cross-sectional views. (G) Schematic of gut-on-a-chip bowel prep infection model. Chips were exposed to vehicle (350 mOsm/kg) or bowel prep (600 mOsm/kg) treatment at high flow rate (600 μL/h), infected statically with *Salmonella* Typhimurium for 30 minutes, then exposed to fluid flow (120 μL/h) until 24 hours post infection. (H) Apparent permeability (Papp) of the gut epithelial layer on the gut-on-a-chip, as measured by FITC-dextran permeability assay 4 hours after exposure to *Salmonella* Typhimurium (Veh *n*=3, Bowel Prep *n*=3). Each dot corresponds to one chip, with mean Papp for a treatment group/time period shown as black bars. (I) Mean *Salmonella* Typhimurium translocation from the luminal side (top channel) to pass the endothelial layer (bottom channel) 24 hours post-pathogen exposure (Veh *n* = 3, Bowel Prep *n* = 3). Both the log scale (left) and the linear scale (right) are shown. Each dot corresponds to one chip, with mean CFU/mL shown as black bars. **Statistics:** For all comparisons, statistical significance was assessed using the Wilcoxon ranked-sum test. p > 0.05; ns (not significant, not shown), p < 0.05; *, p < 0.01; **, p < 0.001; ***, p < 0.0001; ****. **Abbreviation:** LoD, limit of detection.

**Supplemental Figure 7.**
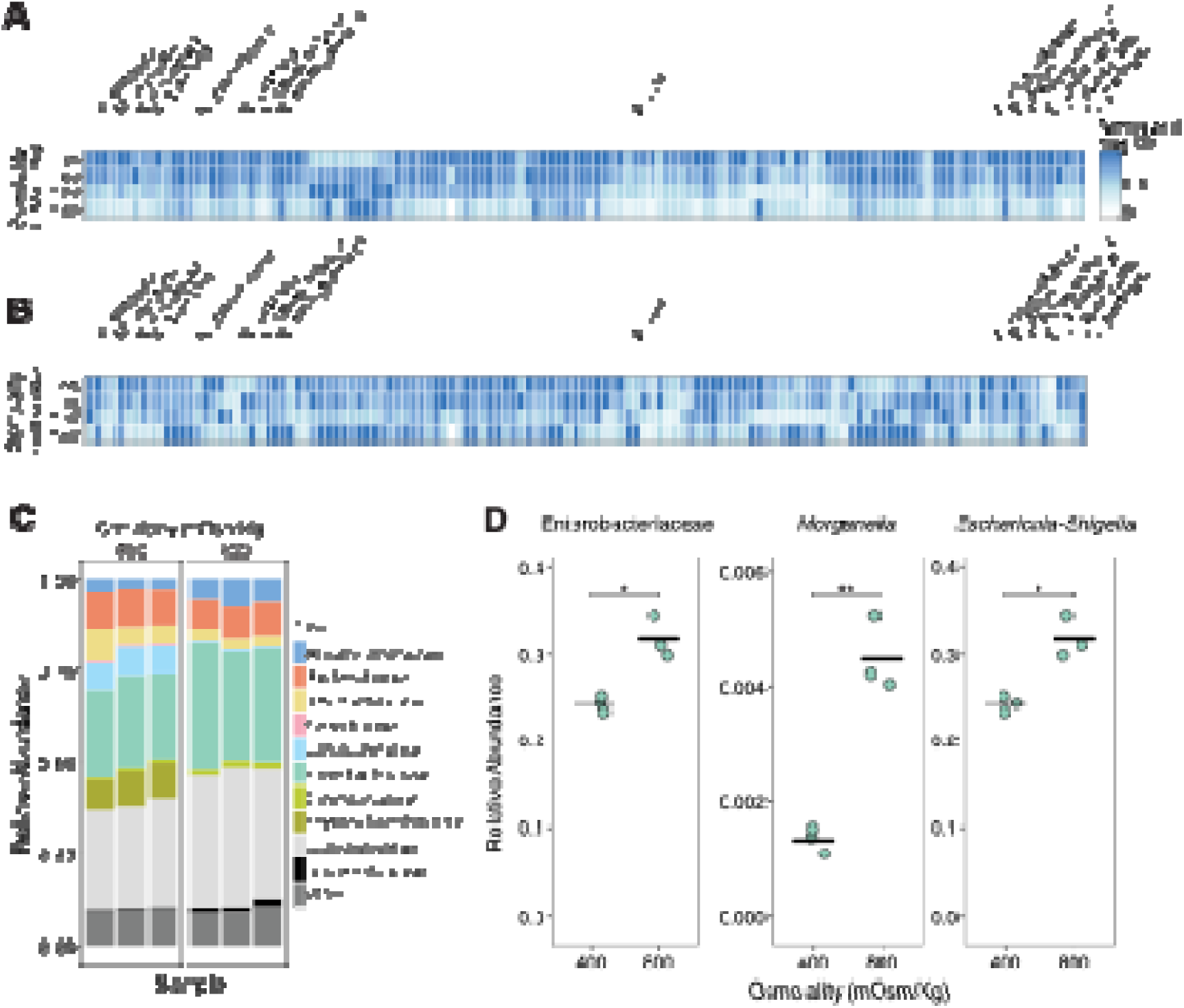
Potential pathobionts from patients with inflammatory bowel disease are resilient to osmotic perturbation *in vitro* and can translocate and persist in the mesenteric lymph nodes of germ-free mice after bowel prep. (A) Growth of 130 potential pathobionts isolated from patients with ulcerative colitis and cultured under anaerobic conditions at osmotic stress levels meeting or exceeding those found in the mouse gut after bowel prep (∼750 mOsm/Kg). Normalized carrying capacity was measured at maximum OD600 after bacteria were cultured under anaerobic conditions for 24 hours, and cultures were grown at 400, 800, 1200, and 1800 mOsm/Kg, adjusted with NaCl. (B) Potential pathobionts from (A) cultured in aerobic conditions. (C) 16S rRNA sequence data from a fecal sample obtained from an individual with ulcerative colitis cultured under osmotic stress levels simulating normal (400 mOsm/Kg) and (PEG)-treated (800 mOsm/Kg) conditions. (D) Relative abundance of the Enterobacteriaceae family and the genera *Morganella* and *Escherichia-Shigella* from (B) (each osmolality *n*=3). Differences between treatment groups were analyzed with t-tests. p > 0.05; ns (not significant, not shown), p < 0.05; *, p < 0.01; **, p < 0.001; ***, p < 0.0001; ****. **Abbreviation:** LoD, limit of detection.

**Supplemental Figure 8.**
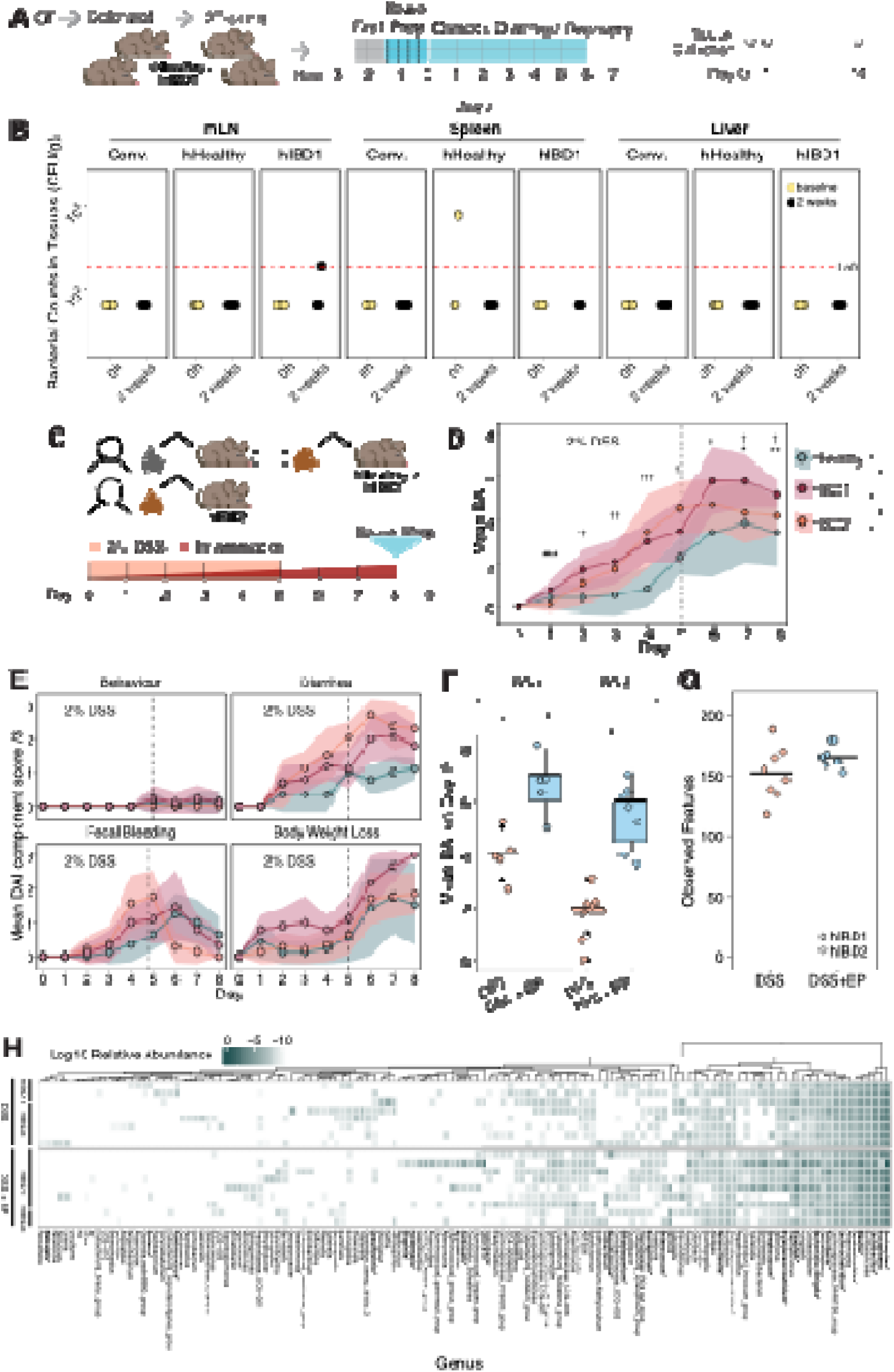
Germ-free mice humanized with microbiota from patients with inflammatory bowel disease experience greater chemically-induced colitis disease than those humanized with a healthy microbiota, in a microbiota-dependent manner. (A) Schematic of germ-free (GF) C57BL/6J mice colonized with a fecal sample from a patient with ulcerative colitis (hIBD1) or without the condition (hHealthy) and then bred for experimental use. Mouse offspring 8-12 weeks of age were bowel prepped and sampled 2 weeks post-bowel prep. (B) Anaerobic bacterial translocation measured in extraintestinal organs, mesenteric lymph nodes (mLN), spleen, and liver, at baseline and 2 weeks post-bowel prep in conventional (Conv.) mice, hHealthy mice, and hIBD mice after bowel prep (Conv. each tissue: baseline *n*=5, 2 weeks *n*=7. hHealthy each tissue: baseline *n*=4, 2 weeks *n*=5. hIBD1 mLN: baseline *n*=4, 2 weeks *n*=4. hIBD Spleen & Liver: baseline *n*=5, 2 weeks *n*=4.). (C) Schematic depicting humanisation of GF CBL57/6J mice with fecal samples from mice previously humanized using healthy (hHealthy) or ulcerative colitis (hIBD1) patient samples, or with fecal samples directly from a patient with ulcerative colitis (hIBD2). Mice were treated with 2% dextran sodium sulphate (DSS) in drinking water for 5 days to stimulate a colitis flare-up. (D) Mean disease activity index (DAI) was assessed daily within the three microbiota. Statistical comparisons between hIBD1 & hIBD2, and hIBD1 & hHealthy, are denoted by * and † respectively. (E) Detailed breakdown of the DAI components show that colitis disease symptoms are dependent on mouse microbiota. Statistics not shown. (F) On day 9, 24 hours after bowel prep, both hIBD microbiota groups show an increase in DAI compared to mice of the same microbiota that did not receive the treatment. (G) Stool 16S rRNA sequencing on day 9. (H) 16S rRNA sequencing of bacteria translocated to the mesenteric lymph nodes of mice from (C-G). Pathobionts associated with IBD^6,48–62^ are bolded. **Statistics:** For all comparisons, statistical significance was assessed using the Wilcoxon ranked-sum test. p > 0.05; ns (not significant, not shown), p < 0.05; *, p < 0.01; **, p < 0.001; ***, p < 0.0001; ****.

**Supplemental Data Table S1. Mouse cecal RNA-seq data and statistics, refers to Fig. S1G-I. Host cecal tissue transcriptomics were performed at Baseline and 6 hours post-bowel prep and analyzed as referred to in the Methods. Description of sheets and columns can be found below.**

**Sheet "rna-seq-raw_counts":** This table summarizes the raw gene expression counts from the extracted RNA.

- CC_1_S59ReadsPerGene.out.tab- CC_6_S64ReadsPerGene.out.tab: Raw gene expression counts under baseline conditions for mice 1–6.
- CC_7_S65ReadsPerGene.out.tab- CC_12_S70ReadsPerGene.out.tab: Raw gene expression counts at 6 hours post-treatment for mice 1–6.

**Sheet "DESeq_lfc_Z":** This table summarizes the results table generated from the DESeq2 analyses as outlined in the methods, and the Z-scores across multiple conditions and time points. The Z-score is the number of standard deviations away from the mean of the expression levels of a gene across all the samples.

- Gene: The official symbol for the gene.
- Description: A brief description of the gene’s function or associated protein.
- Baseline_x: The average normalized counts, normalized by sequencing depth across all samples for mice1-6 under baseline conditions.
- 6hr_x: The average normalized counts, normalized by sequencing depth across all samples for mice 1-6 at 6 hours post treatment.
- Log2FoldChange_x: indicates how much the gene expression changed between the samples collected at 6 hours post-treatment and the samples collected at baseline. This value is reported on a log2 scale.
- lfcSE_x: The standard error estimate for the log2 fold change estimate for the samples collected at 6 hours post-treatments
- pvalue_x: P-value of the test for the gene for the samples collected 6 hours post-treatment
- padj_x: Adjusted P-value for multiple testing for the gene for the samples collected 6 hours post-treatment
- mean_Baseline, mean_6hr: Mean normalized counts for each respective condition, averaged across all replicates.
- Z_Baseline_x: Z-scores representing gene expression levels under baseline conditions for mice 1–6.
- Z_6hr_x: Z-scores for gene expression at 6 hours post-treatment for mice 1–6.
- Z_Baseline_mean, Z_6h_mean: Mean Z-scores for each respective condition, averaged across all replicates.

**Supplemental Data Table S2. Papp calculations, refers to Fig. S6H. Samples from all channels were collected at baseline, 4 hours post infection, and 24 hours post infection and analyzed as referred to in Methods. Description of the sheets can be found below.**

**Sheet “Papp_Calculator”:** This sheet is the template of the Emulate Papp calculator^95^ used in other sheets.

**Sheet “baseline”:** This sheet shows the Papp calculations for samples collected at baseline.

**Sheet “4hpi”:** This sheet shows the Papp calculations for samples collected at 4 hours post infection.

**Sheet “24hpi”:** This sheet shows the Papp calculations for samples collected at 24 hours post infection.

**Supplemental Data Table S3. Outputs from GSEA analysis, refers to Fig. 5C–D. Sheet “GSEA Analysis”**

- ID: Gene Ontology (GO) identifier, a unique accession for a biological process.
- Description: Plain-text name of the GO term associated with the ID.
- setSize: Number of genes from the dataset that fall within the given GO term gene set.
- enrichmentScore: Raw enrichment score (ES) reflecting the degree to which genes in a GO term are overrepresented at the top or bottom of the ranked gene list. Positive ES means enrichment in upregulated genes; negative ES means enrichment in downregulated genes.
- NES: Normalized enrichment score (NES) is the enrichment score normalized for gene set size, allowing comparisons across gene sets of different sizes.
- pvalue: Nominal p-value assessing significance of the observed enrichment score by permutation testing.
- p.adjust: Multiple hypothesis testing–adjusted p-value (via Benjamini–Hochberg correction).
- qvalue: Estimated false discovery rate (FDR) for the enrichment result.
- rank: Position in the ranked gene list where the enrichment score was maximal (peak enrichment).
- leading_edge: Subset of genes that contribute most strongly to the enrichment signal. Reported as percentages of “tags” (genes in the set contributing), “list” (portion of ranked list contributing), and “signal” (relative strength of enrichment).
- core_enrichment: Specific genes from the dataset that belong to the leading edge subset, reported as Entrez Gene IDs.

## CRediT authorship contribution statement

CAC: Conceptualization, Methodology, Formal Analysis, Investigation, Data Curation, Writing – original draft, Writing - Review & Editing, Visualization

IP: Conceptualization, Methodology, Formal Analysis, Investigation, Data Curation, Writing – original draft, Writing – Review & Editing, Visualization

BDD: Conceptualization, Methodology, Formal Analysis, Investigation, Data Curation, Writing – original draft, Writing – Review & Editing, Visualization

GM: Methodology, Formal Analysis, Investigation, Writing - Review & Editing, Visualization

AS: Investigation, Formal Analysis

CS: Investigation

JH: Formal Analysis

AP: Investigation

DT: Investigation

DMP: Investigation

TF: Investigation

KN: Conceptualization, Methodology, Investigation, Writing - Review & Editing, Supervision

SRS: Conceptualization, Resources, Supervision

MS: Resources, Supervision

BAV: Conceptualization, Resources

CT: Conceptualization, Resources, Writing – original draft, Writing– review & editing, Formal Analysis, Supervision, Project administration, Funding acquisition.

## Methods

### Animal Handling and Ethics

All animal experiments were conducted in accordance with the ethical guidelines of the University of British Columbia’s (UBC) animal care procedures, following protocol number A23-0115 approved by the Animal Care Committee. C57BL/6J male and female mice between 8-12 weeks of age were used and were provided with an autoclaved standard diet (Purina LabDiet 5K67). As we did not observe sex differences, male and female mice were analyzed together in all experiments. Experimental groups that underwent oral gavage were monitored at 1 hour and 24 hours post-procedure. Daily health checks for mortality after infection were performed, and mice were euthanized if they lost more than 20% of their body weight or displayed signs of distress. Euthanasia at indicated tissue-collection time points was performed using carbon dioxide asphyxiation followed by cervical dislocation.

### Bowel Prep Mouse Model

Bowel prep solution was prepared by dissolving 42.5% (w/v) polyethylene glycol (PEG) (commercially branded Restoralax) in ddH_2_O and filter sterilized. Food was withdrawn from all cages 1 hour before the bowel prep procedure. Mice were orally gavaged with 200 μL of bowel prep solution every 20 minutes a total of 4 times. Vehicle-treated mice were orally gavaged with 200 μL of water following the same dosing time as bowel prep-treated mice. Food and water were readministered after the procedure.

This concentration and delivery protocol were selected to reflect PEG dosing used in human bowel prep regimens, which typically involve administration of up to 420 g of PEG over a short time frame, corresponding to an estimated 4.9–10.5 g PEG/kg body weight depending on patient size. In our model, mice received a total of ∼0.34 g PEG (administered as 200 μL every 20 minutes for four doses), corresponding to ∼7.2–13.6 g PEG/kg, a dose range physiologically comparable to human protocols when adjusted for mouse body weight and intestinal clearance. To account for potential hydration differences introduced by the vehicle, mice were monitored for signs of dehydration and provided with ad libitum access to water immediately following the procedure.

We additionally evaluated the impact of high dose osmotic laxatives on the gut environment and the gut microbiota at 6, 24, 48, and 72 hours post-bowel prep compared to untreated controls sacrificed at the 6-hour timepoint (baseline).

To investigate the effects of bowel prep on the gut environment, we characterized a murine bowel prep method previously used to facilitate the transplantation of human microbes into mice^96^. Mice with a conventional microbiota were fasted and then orally gavaged 4 times with PEG dissolved in water (bowel prep), while controls received water as the vehicle alone (Fig. 1A). Mice were euthanized 6 hours post-bowel prep, and intestinal tissues and contents were collected. Bowel prep-treated mice had an average cecal osmolality that was 1.8-fold higher than vehicle-treated controls (725 vs. 437 mOsm/Kg, p < 0.001) (Fig. 1B) and a higher cecal mass, indicative of osmotic diarrhea and increased luminal contents during bowel prep (Fig. S1A).

### Cecal Osmolality and pH Measurements

Mice were sacrificed, and cecal contents were collected and placed on ice. The osmolality of the cecal contents was directly measured using an Advanced Instruments Osmo1 Single-Sample Micro-Osmometer (Fisher Scientific). pH was measured with a calibrated micro pH probe (Orion PerpHecT ROSS Combination pH Micro Electrode).

### Tissue Collection

Sterile tools were used to collect the lower lobe of the liver, mesenteric lymph nodes, and the spleen. The tissues were placed in pre-weighed sterile 2 mL tubes containing 200 μL of PBS and were put on ice. Cecal contents were collected, and half of the contents were flash-frozen on dry ice, while the other half was placed on ice for downstream analysis. The ileum, cecum, and colon tissues were collected in histology cassettes and immediately fixed in methacarn solution (60% dry methanol [Fisher Chemical], 30% chloroform [Fisher Chemical], 10% glacial acetic acid [Fisher Chemical]) for 7 to 14 days. The samples were then washed twice in methanol for 30 minutes, twice in 100% ethanol (Sigma) for 20 minutes, and twice in xylenes (Fisher Chemical) for 15 minutes. The tissues were coated and incubated in paraffin at 60°C for 2 hours and then dried at room temperature. Paraffin blocks were cut into 4 μm sections and mounted on slides by the British Columbia Children’s Hospital Research Institute’s Histology Core^97^.

### Lectin Staining of the GIT

Paraffin was removed from sectioned slides through incubation at 60°C for 10 minutes, followed by 2x 10-minute incubations in pre-warmed xylenes at 60°C. Slides were then incubated in 99.5% ethanol for 5 minutes, then left to dry and circled with a PAP (liquid blocker) pen (Fisher Scientific)^97^. DAPI (10 μg/mL, Sigma Aldrich), UEA-1 (Fluorescein-labeled Ulex Europaeus Agglutinin I, 40 μg/mL, Vector Laboratories), and WGA (Rhodamine Red-X-labeled Wheat Germ Agglutinin, 30 μg/mL, Vector Laboratories) were applied to fully cover the samples (approximately 250 μL) and incubated in the dark at 4°C for 45 minutes (Table 1). The slides were then washed in PBS 3 times. Sections were left to dry for 5 minutes, and ProLong Gold Antifade Mountant, (Invitrogen), was applied, followed by a #1.5 glass coverslip^97^. Images were collected using a Zeiss LSM 900 confocal microscope at 100x (**Fig. 1, S1**) or 20x (**Fig. 3, S3**) magnification with the ZEN 2020 software. Mucus thickness was quantified using the analysis platform BacSpace^98^.

### Confocal Image Selection and Mucus Quantification

Confocal images of longitudinal sections of the cecal tip and distal colon were collected for 5-8 mice per treatment group. For each tissue, three representative sites with ≥500 µm continuous epithelium were imaged per mouse. Whenever possible we used sections that contained intestinal contents; however, in the case of bowel prep the contents were purposely evacuated by the procedure. These sites were selected for continuous epithelium and the presence of intact intestinal contents, where possible.

The contour defining the boundary of the epithelium and the mucus layer were identified and defined using the BacSpace MatLab software^98^. A mucus layer was defined as having minimum thickness of 5 µm and no non-bacterial gut contents (e.g., large fibre pieces). Using a custom MatLab Script, mucus-covered area was defined as the region(s) between these curves. Percentage mucus coverage was calculated as the total length of the mucus divided by the total length of the epithelium. Average mucus thickness was calculated as the total covered by mucus divided by the total length of the epithelium. Quantitative analyses were performed on each image, and the mean value from the three images was calculated to represent a single data point per mouse. Differences in average thickness between treatment and control groups (Figure 1) or between timepoints post-bowel prep (Figs. 3, S3) were assessed using a one-way ANOVA.

### Hematoxylin & Eosin Tissue Histology

Paraffin was melted from the section slides by heating the slides in coplin jars at 60°C for 10 minutes. Slides were then incubated in the coplin jars in 60 °C xylenes for 3 minutes twice followed by 2 incubations in 100% EtOH for 2 minutes each, 95% EtOH for 2 minutes, and in ddH20 for 2 minutes^99^. Next, the hematoxylin stain was performed by filling the coplin jars with Hematoxylin Solution, Gill’s No. 2 (Sigma Aldrich, GHS216), and incubating for 3 minutes, then in ddH_2_O for 1 minute, differentiator solution (0.3% v/v 10N HCl [Fisher Chemical], 70% v/v EtOH [Sigma] in ddH2O)) for 30 seconds, ddH20 for 1 minute, and blueing reagent (0.2% w/v NaHCO_3_ [Fisher BioReagents], 4.1% w/v MgSO_4_ 6H_2_O [Fisher BioReagents] in ddH2O)) for 1 minute. For the Eosin stain, slides remained in the coplin jar and were filled with 95% EtOH for 1 minute followed by Eosin Y solution (Sigma Aldrich), for 45 seconds. The stain was followed by washes in 95% EtOH for 1 minute, in 100% EtOH for 1 minute 3 times, and in xylenes for 2 minutes twice. Stained slides were mounted using Permount (Fisher Chemical), a xylene-based mounting medium, and allowed to solidify for at least 24 hours^99^. H&E images were acquired using the 3D HISTECH Slide Scanner at 37x (Fig. 1E, 2E) or the ZEISS Digital Slide Scanner Axioscan 7 at 20x (Fig. 4C,6D).

### *Salmonella* Typhimurium Infection Pathology Scoring

H&E-stained sections of the cecum and distal colon were assessed for pathology in the lumen, epithelium, mucosa, and submucosa of the gut. Luminal pathology was scored based on the presence of necrotic epithelial cells and polymorphonuclear neutrophils (PMNs) (0 = none, 1 = scant, 2 = moderate, 3 = dense). The epithelium was scored for desquamation (0 = no change, 1 = limited shedding, 2 = moderate shedding per lesion), regenerative change (0 = none, 1 = mild, 2 = moderate, 3 = severe), ulceration (1 = epithelial ulceration), and PMNs in the epithelium (1 = PMNs present). The mucosa was scored based on crypt abscesses (0 = none, 1 = mild, 2 = moderate, 3 = severe) and the presence of mucin plugs and granulation tissue (1 = present). Lastly, the submucosa was scored for edema, mononuclear cell infiltration, and PMN infiltration (0 = no change, 1 = mild, 2 = moderate, 3 = severe). Scores for each location across the image were summed together, with the maximum pathological score being 24 per image. Two images of 1000 µm continuous mucosa were taken for each mouse, and were scored by two blinded scorers. Scores were averaged for each mouse.

### Mucosa Thickness Measurements

20X images of H&E-stained tissues were exported in TIFF format from SlideViewer. TIFF images were converted from RGB to 8-bit format in FIJI, and pixel values were inverted so that tissues appeared light against a dark background. BacSpace was used to create two masks; one mask outlined the submucosal edge of the mucosa, and the other mask outlined the luminal edge of the epithelium. Both masks were applied to the original image so that only the mucosa and epithelium were non-zero values. BacSpace was then used on the masked images to straighten the image relative to the luminal edge of the epithelium. The luminal edge of the epithelium in the straightened image was detected as the greatest decrease in signal along the x-axis of the straightened image at each point along the epithelium. The mean thickness was quantified for each image, and the mean thicknesses of two images from the cecum and two images from the distal colon from three different mice per treatment (total = 6 images per treatment per organ) were compared using a Wilcoxon rank test.

### Quantification of Gut Permeability

Intestinal permeability was assessed in mice following bowel prep using a FITC-Dextran assay^100,101^. Mice with a conventional microbiome received bowel prep (*n* =12) or a vehicle control (*n* = 12), then continued to be fasted for an additional 2 hours before receiving 150 µL of 80 mg/mL fluorescein isothiocyanate (FITC)–Dextran (4 kDa, Sigma-Aldrich) in PBS by oral gavage. Food and water were readministered after the FITC-Dextran gavage. Mice were sacrificed 4 hours following FITC-Dextran administration, and serum was collected by cardiac puncture. The fluorescence of FITC in serum was measured in duplicate using emission wavelength of 485 nm and emission wavelength of 535 nm using a Biotek Synergy H1 plate reader. A standard curve was generated using serial dilutions of FITC-Dextran, and serum from untreated mice was used to account for background fluorescence.

### Quantification of Bacterial Levels in Feces

1 μL of feces was diluted in 200 μL of sterile PBS. The solution was serially diluted at 1:10 down a 96-well plate. Next, 5 μL fecal dilutions were spot-plated on both LB-streptomycin agar plates and Columbia Blood agar plates. Plates were incubated aerobically overnight at 37 °C. Single colonies were counted from the highest possible dilution and back calculated to determine the absolute microbial abundance.

### DNA Extraction, Library Preparation, 16S rRNA Sequencing and Analysis

DNA was extracted from fecal pellets and cecal contents using the DNeasy 96 PowerSoil Pro QIAcube HT Kit (Qiagen Inc., Valencia, CA) according to the manufacturer’s instructions.

For all 16S rRNA runs except for the *in vitro* IBD (**Fig. S7A-D**), 16S rRNA library preparation was conducted at the Biofactorial High-Throughput Biology (Bio!) Facility at the University of British Columbia. Amplification of the V4 region was performed with 515F/926R primers (515F, 5’-GTGYCAGCMGCCGCGGTAA-3’; 926R, 5’-CCGYCAATTYMTTTRAGTTT-3’). Pooled libraries were then submitted to the Bio! facility, where sequencing was performed on the Illumina MiSeq^TM^ platform with v2 2 x 300 bp paired-end read chemistry. For the *in vitro* IBD samples, 16S library preparation was performed by Gut4Health as previously described^102^. Amplicons of the V4 region of the 16S rRNA were generated using KAPA HiFi HotStart Real-time PCR Master Mix (Roche) and barcode primers 515F: GTGYCAGCMGCCGCGGTAA and 806R: GGACTACNVGGGTWTCTAAT. Purified PCR libraries were normalized and pooled with the SequalPrepTM normalization plate (Applied Biosystems). Library concentrations were confirmed using the QubitTM dsDNA high sensitivity assay kit (Invitrogen) and KAPA Library Quantification Kit (Roche). The purified pooled libraries were then submitted to UBC’s Bioinformatics and Sequencing Consortium (SBC). Paired end read sequencing was carried out either on the Illumina MiSeqTM v3 platform with 2 x 300 bp paired end-read chemistry or the NextSeq 600 cycle P1 with 2 x 301 bp. To ensure DNA quality and quantity, an Agilent high sensitivity DNA kit (Agilent) was employed on an Agilent 2100 Bioanalyzer.

Read quality was assessed by running FASTQC^103^ on the generated FASTQ files. Reads were then imported into QIIME2-2023.9 for subsequent analyses^104^. DADA2 (via q2-dada2) was used to denoise and quality filter the data; then reads were trimmed to remove primer sequences while maintaining mean Phred quality scores >Q30^105^. Using the QIIME classification plugin (q2-feature-classifier), amplicon sequence variants (ASVs) were classified via a naïve Bayes machine-learning taxonomic classifier against the SILVA 138 99% identity reference sequence database^106^. Multiple sequence alignment and phylogenetic tree generation was performed using MAFFT (via q2-alignment) and FastTree2 respectively (via q2-phylogeny)^107,108^. Plotting was conducted using R v4.2.2^109^. Tidyverse^110^ and ggplot2^111^ packages were used for data visualization. The packages phyloseq^112^, ggpubr^113^, and vegan^114^ were used for sample rarefaction, calculation, and visualization of alpha and beta diversity metrics.

### SCFA Extraction from Cecal Contents

SCFAs were extracted from 40 to 100 mg of flash-frozen cecal contents. The samples were homogenized with 0.8 mL of 25% phosphoric acid (LabChem) and centrifuged at 15,000 × g for 10 minutes at 4°C. The supernatant was removed from all samples and centrifuged again. Subsequently, 800 μL of the supernatant was filtered through a 0.45 µm filter (Fisher) and mixed with 0.2 mL of the internal standard solution containing 24.5 mmol/L isocaproic acid in a GC vial (12×32mm, Thermo Scientific). SCFAs were quantified by gas chromatography-mass spectrometry by the AFNS Chromatography Facility at the University of Alberta, as previously described^115^.

### *Salmonella* Typhimurium Culture and Inoculum Preparation

*Salmonella* Typhimurium Δ*flhD* was a gift from the Finlay Lab at UBC and mCherry-*Salmonella* Typhimurium was a gift from the Vallacne Lab at UBC. Glycerol stocks of the naturally streptomycin resistant *Salmonella enterica* serovar Typhimurium SL1344 strains (*Salmonella* Typhimurium [WT] and *Salmonella* Typhimurium Δ*flhD*, mCherry-*Salmonella* Typhimurium) were streaked onto Luria-Bertani Miller (LB) 1.5% agar plates supplemented with streptomycin (100 µg/mL). Following incubation at 37°C for 24 hours without agitation, a single *Salmonella* Typhimurium colony was selected from the agar plate and transferred to 5 mL of LB liquid broth containing 100 µg/mL streptomycin. The culture was then aerobically incubated at 37°C with shaking at 200 rpm for 17 hours. Subsequently, the cultures were serially diluted in PBS to achieve the desired inoculant bacterial counts.

### *Salmonella* Typhimurium Osmolality Growth Measurements

Wildtype *Salmonella* Typhimurium or *Salmonella* Typhimurium Δ*flhD* were diluted 1:100 from overnight LB cultures in fresh LB media adjusted to ∼400, 800, 1200, and 1800 mOsm/Kg using PEG (Restoralax). Growth curves were obtained using a Biotek Synergy H1 plate reader at 37°C for 24 hours aerobically. The plates were shaken orbitally every 15 minutes following the collection of OD600 measurements. Background absorbance subtraction was performed from non-inoculated wells. The growth rates of the strains were determined using a previously published MATLAB package^19^.

### Streptomycin *Salmonella* Typhimurium Infection Model

Conventional mice were fasted for 4 hours before being treated with 100 μL of 200 mg/mL streptomycin (Sigma-Aldrich, S6501-50G) by oral gavage^32^. 24 hours later, mice were fasted for 3 hours and infected by oral gavage with 100 μL of 10^6^ CFU *Salmonella* Typhimurium. Fecal samples were collected daily for three days after *Salmonella* Typhimurium infection and spot-plated on streptomycin-LB agar plates.

### Bowel Prep *Salmonella* Typhimurium Infection Model

Conventional mice underwent bowel prep as described previously and received oral gavage with 100 μL of 10^6^ CFU of wildtype *Salmonella* Typhimurium or *Salmonella* Typhimurium Δ*flhD* 6 hours after the bowel prep. To assess dose-dependent *Salmonella* Typhimurium colonization, mice were infected with 10^2^, 10^3^, 10^5^, 10^6^, and 10^9^ CFU post-bowel prep. To explore the susceptibility to *Salmonella* Typhimurium infection following bowel prep over time, mice were infected with 10^6^ CFU of *Salmonella* Typhimurium 6, 24, and 48 hours post-bowel prep. Fecal samples were collected daily for three days after *Salmonella* Typhimurium infection and spot-plated on Columbia Blood agar plates (per liter: 35 g Columbia Broth, 5% Sheep Blood, hemin, and vitamin K) to determine total absolute bacterial abundance and streptomycin-LB agar plates to determine absolute abundance of *Salmonella* Typhimurium.

### RNA extraction, Sequencing and Analysis

Cecal tip tissue was collected from conventional CL57/6J mice treated with bowel prep and sacrificed 6 hours later alongside an untreated baseline group. RNA was extracted from cecal tip tissue using the RNeasy Mini Kit (Qiagen) according to the manufacturer’s instructions. The extracted RNA was sent to the Sequencing and Bioinformatics Consortium at UBC, where a sequencing library was prepared using a standard Illumina Stranded mRNA kit (Illumina) to generate 150 bp pair-end reads. Raw reads were assessed for nucleotide quality (terminal bases below PHRED quality score 20 were removed), trimmed to remove adaptor sequences, and merged using Fastp v. 0.23.4^116^. The reads were then aligned to the mouse genome GRCm39 using the STAR 2.7 software^117^ with --quantMode GeneCounts, and the gene counts obtained were imported into R for downstream analyses.

Raw gene counts were analyzed using DESeq2^118^. Volcano plots of genes with adjusted p-value (Benjamin-Hochberg corrected) less than 0.05 and log_2_ fold change greater than 1.5 were plotted using the EnhancedVolcano package^119^. Heatmaps of 40 select genes of interest and the top 50 most variable genes by absolute Z-score were plotted with the pheatmap package^120^. Gene set enrichment analysis (GSEA) was performed using the clusterProfiler package^121^. Genes were ranked by multiplying the sign of log_2_ fold change with the −log_10_(p-value) obtained from the DESeq2 results. Gene symbols were converted to Entrez IDs using the org.Mm.eg.db package^122^ and enrichment analysis was performed using the Gene Onology (GO) biological process database including gene sets between 10 and 400. The normalized enrichment score (NES) of GO terms with a p-value below 0.05 was visualized using the ggplot2 package^111^. In addition, the log_2_ fold change of genes within the leading edge subset of each enriched GO term was extracted and plotted using the ggridges^123^ and upsetR^124^ packages.

### Mammalian Cell Culture

Caco-2 and HT-29 cells were cultured individually in Dulbecco’s Modified Eagle Medium (DMEM) high glucose (Sigma-Aldrich) supplemented with 10% (v/v) heat-inactivated fetal bovine serum (FBS, Corning), 1% (v/v) Penicillin-Streptomycin (Sigma-Aldrich), 1% (v/v) GlutaMAX supplement (Thermo Fisher), and 1% (v/v) non-essential amino acid (NEAA, Sigma-Aldrich) solution in T75 flasks (Corning). Cells were grown in humidified incubators at 37°C and 5% CO_2_. Media was replaced every other day and cells were passaged once they reached 60-70% confluency using 3 mL of TrypLE (Thermo Fisher) in a 1:10 split ratio.

Human Intestinal Microvascular Endothelial Cells (HIMECs) were cultured with complete endothelial cell growth medium (Sigma-Aldrich) and 1% (v/v) Penicillin-Streptomycin (Sigma-Aldrich) in humidified incubators at 37°C and 5% CO_2_. Frozen cells at passage 7 were thawed a week before experiment and seeded into a T75 flask (Corning) coated with 5 mL of Attachment Factor (Cell Systems). Media was changed every other day and cells were passaged once at 80% confluency using 3 mL of TrypLE by seeding 1 x 10^5^ cells into an Attachment Factor-coated T75 flask.

### HT-29 Monolayer PEG Exposure, Invasion, and Gentamicin Protection Models

Circular glass coverslips (Fisherbrand) in a 24-well plate (Corning) were coated with 100 µL of Bovine Collagen I (Corning) diluted 1:60 in 0.01 N hydrochloric acid (Fisher Chemical). HT-29 cells between passages 18-25 were detached with TrypLE (Thermo Fisher) and resuspended to 2 x 10^6^ cells/mL, then 100 µL of cells were seeded to each coated coverslip along with 400 µL of supplemented DMEM media. Cells were cultured for four days before *Salmonella* Typhimurium infection, and media were changed every second day. The day before infection, media were exchanged with antibiotic-free supplemented DMEM.

For PEG dosage experiments, basal DMEM media adjusted to 600, 900, or 1200 mOsm/kg with PEG was added to each well at 24 hours, 3 hours, and 1 hour prior to infection. Monolayers were infected with late-log phase *Salmonella* Typhimurium at a multiplicity of infection (MOI) of 50. After 30 minutes, all wells were treated with 100 µL/mL gentamicin (Sigma Aldrich) in basal DMEM for 1 hour. To obtain counts for intracellular invasion, monolayers were lysed with 1% Triton X-100 (MilliporeSigma) and 1:10 serial dilutions were plated on LB-strep plates.

For flow cytometry and microscopy experiments, basal DMEM media adjusted to 600 mOsm/kg with PEG was added to each well 3 hours before infection. Monolayers were infected with late-log phase mCherry-expressing *Salmonella* Typhimurium^38^ at an MOI of 50. After 1 hour, all wells were treated with 10 µL/mL gentamicin in basal DMEM for 2 hours.

### Flow cytometry of HT-29 Monolayers

Following infection endpoint, cells were detached with TrypLE (Thermo Fisher) and pelleted at 5,000 x g for 5 minutes, then fixed in 2% paraformaldehyde and resuspended in PBS to be analyzed on a CytoFLEX LX flow cytometer. mCherry signal was excited with the 561 nm yellow laser and detected using a 610/20 nm bandpass filter. Cells and singlets were gated, and at least 25,000 events were recorded from the singlets gate for each sample at medium speed (30 µL/minute). FCS files were imported into FlowJo (v10.10.0) and an mCherry-positive gate was defined using negative control samples, with thresholds set to include <1% of events from the control populations. All events within the singlets gate for each sample were exported and visualized in R using the ggplot2 package^111^.

### Staining and imaging of HT-29 Monolayers

Monolayers at infection endpoint were fixed with 4% paraformaldehyde for 15 minutes. Cells were permeabilized with 0.2% Triton X-100 and blocked with 5% normal donkey serum (NDS) and 1% bovine serum albumin (BSA) in PBS. Cells were stained with DAPI (Sigma Aldrich) and Alexa-Fluor 488-conjugated phalloidin (Fisher) and mounted to slides using Vectashield (BioLynx). Cells were imaged using a Zeiss LSM 900 confocal microscope and analyzed using ImageJ Fiji (v. 2.16.0/1.54p). 50 Z-stacks with 0.5 µm intervals were taken of each image to ensure coverage of the entire cell, and infected cells and mCherry-*Salmonella* Typhimurium was counted from max intensity projections of images. Four fields of view (FOVs) from each of three technical replicates of each condition were imaged and analyzed, across three independent experiments.

### Gut-on-a-Chip Culture

S1 chips (Emulate) were surface activated according to the manufacturer’s protocol. Chips were then coated overnight in humidified incubators at 37°C and 5% CO_2_ with 200 μg/mL Collagen IV (Sigma-Aldrich) and 100 μg/mL Matrigel (Corning) in the apical channel, and 200 μg/mL Collagen IV and 30 μg/mL Fibronectin (Corning) in the basolateral channel, prepared in sterile cell-culture grade water. The following day, HIMECs at passage 8 were detached with TrypLE then resuspended to 8 x 10^6^ cells/mL and 15 μL was seeded to the basolateral channel. Chips were flipped upside down and incubated for 1 hour to allow HIMECs to adhere. After flipping chips back, 4 parts of Caco-2 at passage 17 and 1 part of HT-29 at passage 16 detached with TrypLE were combined and resuspended to 7.5 x 10^6^ cells/mL, and 35 μL was seeded to the apical channel. Cells were allowed to adhere overnight in the incubator, before connecting chips to continuous flow (60 μL/h) and mechanical deformation (10%, 0.15 Hz) and grown for 3 additional days before switching to antibiotic-free DMEM high glucose with supplements in both channels.

### Chip Bowel Prep and *Salmonella* Typhimurium Translocation Model

Bowel prep solution was prepared by dissolving 0.05 M polyethylene glycol (PEG) (commercially branded Restoralax) in DMEM high glucose or Hanks’ Balanced Salt Solution (HBSS, Thermo Fisher). After two days of culture in antibiotic-free media, chips were treated with DMEM high glucose (∼350 mOsm/kg) or bowel prep solution made with DMEM high glucose (∼630 mOsm/kg) for 30 min at a 600 μL/h flow rate. Chips were disconnected from flow, and channel inlets and outlets were plugged with 200 μL filter tips as 50 μL of 10^6^ CFU/mL *Salmonella* Typhimurium was seeded into the apical channel in HBSS or bowel prep solution made in HBSS, depending on the condition. Bacteria infected statically for 1 hour before filter tips were removed and chips were connected back to flow (120 μL/h) and mechanical deformation (10%, 0.15 Hz) in the incubator, with antibiotic-free DMEM high glucose with supplements. Media from all channels was sampled and spot-plated on LB-streptomycin agar plates at 4 hours and 24 hours post infection. After 24 hours, chips were disconnected then all channels were washed thrice with PBS (Gibco), fixed in 4% paraformaldehyde (Electron Microscopy Sciences, 15710-S) diluted with DPBS (Gibco).

### Paracellular Permeability Analysis

To assess paracellular permeability, 50 µg/ml of 3-5 kDa Fluorescein isothiocyanate-dextran (FITC-dextran, Sigma-Aldrich) was added to the apical channel media on the day before *Salmonella* Typhimurium infection. Chips were run under a 600 μL/h flow rate for 5 min to flush out existing media, then run at a 100 μL/h flow rate for 3 hours before samples from all channels were collected. Media was sampled prior to *Salmonella* Typhimurium infection to establish levels at baseline, after 4 hours of infection, and after 24 hours. The fluorescence intensities (490 nm/520 nm) of the top and bottom channel effluents were measured using a Synergy H1 plate reader (Biotek) and FITC-dextran concentration was estimated by plotting a log-log standard curve. The apical to basolateral flux of FITC-dextran was calculated according to the following equation using the Emulate Apparent Permeability Calculator EC004v1.0^95^:

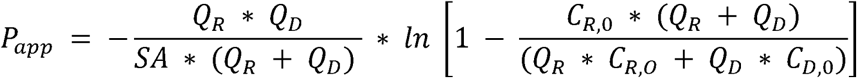

where *P_app_* is the apparent permeability (cm/s); *SA* is the surface area of the sections of the channels that overlap (0.17 cm^2^); Q_R_ and Q_D_ are the fluid flow rates in dosing and receiving channels, respectively (cm^3^/s); and C_R,0_ & C_D,0_ are the recovered concentrations in the dosing and receiving channels, respectively.

### Quantification of Bacterial Colonization in Extraintestinal Tissues

Liver, mesenteric lymph nodes, and spleen were collected in pre-weighed sterile tubes filled with 150 µL of PBS and weighed for final tissue mass. Tissues were homogenized at 300 Hz for 6 minutes using the TissueLyser II (QIAGEN). Homogenized samples were serially diluted at 1:10 in PBS, and all dilutions were spot-plated on both LB-streptomycin agar plates and Columbia Blood agar plates. Plates were incubated aerobically (in the case of *Salmonella* Typhimurium infection) or anaerobically (in the case of hIBD pathobiont translocation) overnight at 37°C. Single colonies were counted and back calculated to determine the absolute abundance of bacterial translocation to extraintestinal organs (liver, mesenteric lymph nodes, and spleen). Excess homogenized tissue was frozen at −80°C for downstream 16S rRNA sequencing.

### Pathobiont Growth Measurements

1 µL of glycerol stocks of each of the 130 pathobionts isolated from UC patients were streaked on LB agar plates. A single colony was inoculated into 5 mL of liquid LB-streptomycin broth and grown for 17 hours overnight aerobically and shaking at 200 rpm. All strains were subcultured 1:15 into LB broth in 2 mL 96 well plates, using a pipetting robot (INTEGRA VIAFLO384) and grown for 2 hours at 37°C^19^. Strains were then inoculated at a ratio of 1:75 into LB media with pH levels of 4.0, 5.5, 6.9, and 8.0, and osmolality levels of 400, 800, 1200, and 1800 mOsm/Kg, adjusted with PEG. The experiments were conducted in technical triplicates using a 384-well plate.

Pathobiont bacterial growth curves were obtained using a Biotek Synergy H1 plate reader at 37°C for 24 hours either inside an anaerobic chamber (Coy Laboratories) with an atmosphere of 5% CO_2_, 5% H_2_, and 90% N2 (Linde Canada) or in aerobic conditions in parallel. The plates were shaken orbitally every 15 minutes following the collection of OD600 measurements. Background absorbance subtraction was performed from non-inoculated wells. The growth rates of the strains were determined using a previously published MATLAB package^19^.

### Human Samples

All studies were approved by the Stanford University Institutional Review Board (IRB). Fecal samples were collected from patients with confirmed ulcerative colitis (UC) and from healthy controls. Stool samples were collected by patients at home in BIOME-Preserve tubes according to manufacturer instructions and returned to the research team within 24 hours, then snap-frozen with dry ice and stored at −80°C until use. The fecal inocula for gavage were prepared in an anaerobic chamber, where 5 g of feces from stool samples was resuspended in 10 mL of sterile and pre-reduced PBS and centrifuged at 4000 RPM for 5 minutes.

### *In vitro* hIBD Microbiota Growth

2 g of human stool inocula were grown from the frozen fecal samples in 5 mL of MEGA medium^19,125^ aerobically overnight and subcultured 1:200 in fresh medium at baseline or adjusted to 800 mOsm/Kg with PEG. 16S rRNA sequencing was performed as described above (DNA Extraction, Library Preparation, and 16S rRNA Sequencing).

### Human Microbiota-Associated Mice

C57BL/6J germ-free mice were inoculated with 200 µL of prepped fecal samples from human donors at 6 weeks of age by oral gavage (ulcerative colitis, hIBD or healthy control, hHealthy, respectively) and maintained in isocages (Tecniplast) for 6 weeks to allow the microbiome to settle. Mice were used directly in experiments (**Fig, 6A-H, S8C-H**) or bred for offspring that were used (hIBD1**, Fig. S8A,B**). Stool was sampled approximately daily.

### Colitis Disease Activity Index

Disease Activity Index (DAI) was measured daily based on the body weight loss, stool consistency and blood in stools using a previously validated approach^42–44^. A higher DAI indicates worse disease state and inflammation. Body weight loss was scored as follows: score 0 for no body weight loss, score 1 for 0-5% weight loss, score 2 for 5-10% weight loss, and score 3 for >10% weight loss. Stool consistency was scored as follows: score 0 for normal faecal pellet, score 1 for soft but adherent in pellet shape, score 2 for loose stool, and score 3 for diarrhoea. Fecal bleeding was scored as follows: score 0 for hemoccult negative, score 1 for hemoccult positive, score 2 for hemoccult positive with visual pellet bleeding, and score 3 for gross visual pellet and rectal bleeding. Behaviour was scored as follows: score 1 for piloerect and some lethargy, score 2 for piloerect and lethargic with little interest in environment, score 3 for high lethargy with no interest in or response to environment or cage mates.

### Mouse IBD DSS Flare-Up Model

Humanized IBD mice received 2% Dextran Sodium Sulphate (DSS, ThermoFisher) in drinking water for 5 days to induce an acute colitis flare-up. Two days after DSS ended mice received bowel prep, or no treatment as a negative control. No vehicle control was used, as DSS treatment itself induces dehydration. This approach better reflects the clinical context of IBD patients recovering from colitis with or without bowel prep and avoids artificially altering hydration dynamics. Mice were sacrificed 24 hours after bowel prep, and tissues and cecal contents were collected. Feces were collected at day 0 (start of DSS), day 8 (bowel prep) and day 9 (day of euthanasia). DAI was assessed daily as described above (Colitis Disease Activity Index). Tissue was homogenized and plated as described above (Quantification of Bacterial Colonization in Extraintestinal Tissues).

### Colitis Pathology Scoring

H&E-stained sections of the cecal tip and distal colon (Hematoxylin & Eosin Tissue Histology) were assessed for pathology following previously published protocols for DSS-induced colitis^47^. The following features of colitis were each scored on a scale of 0-3: immune cell infiltration extent and severity, goblet cell loss, crypt density, crypt hyperplasia, crypt abscesses, muscle thickening, and ulceration. The scores were summed, with the maximum pathological score being 24 per image. Two images of 1000 µm continuous mucosa were taken for each mouse, and were scored independently by two blinded scorers. Scores were then averaged for each mouse. Representative images shown in-text were selected for scoring close to the average score for that treatment group.

### Statistics in Figures

For plating experiments where no CFUs were detected, the values were plotted at ½ the limit of detection (LoD) in log scale. No outlier detection was performed prior to statistical analysis and plotting. p > 0.05; ns (not significant, not shown), p < 0.05; *, p < 0.01; **, p < 0.001; ***, p < 0.0001; ****.

### Key resources table

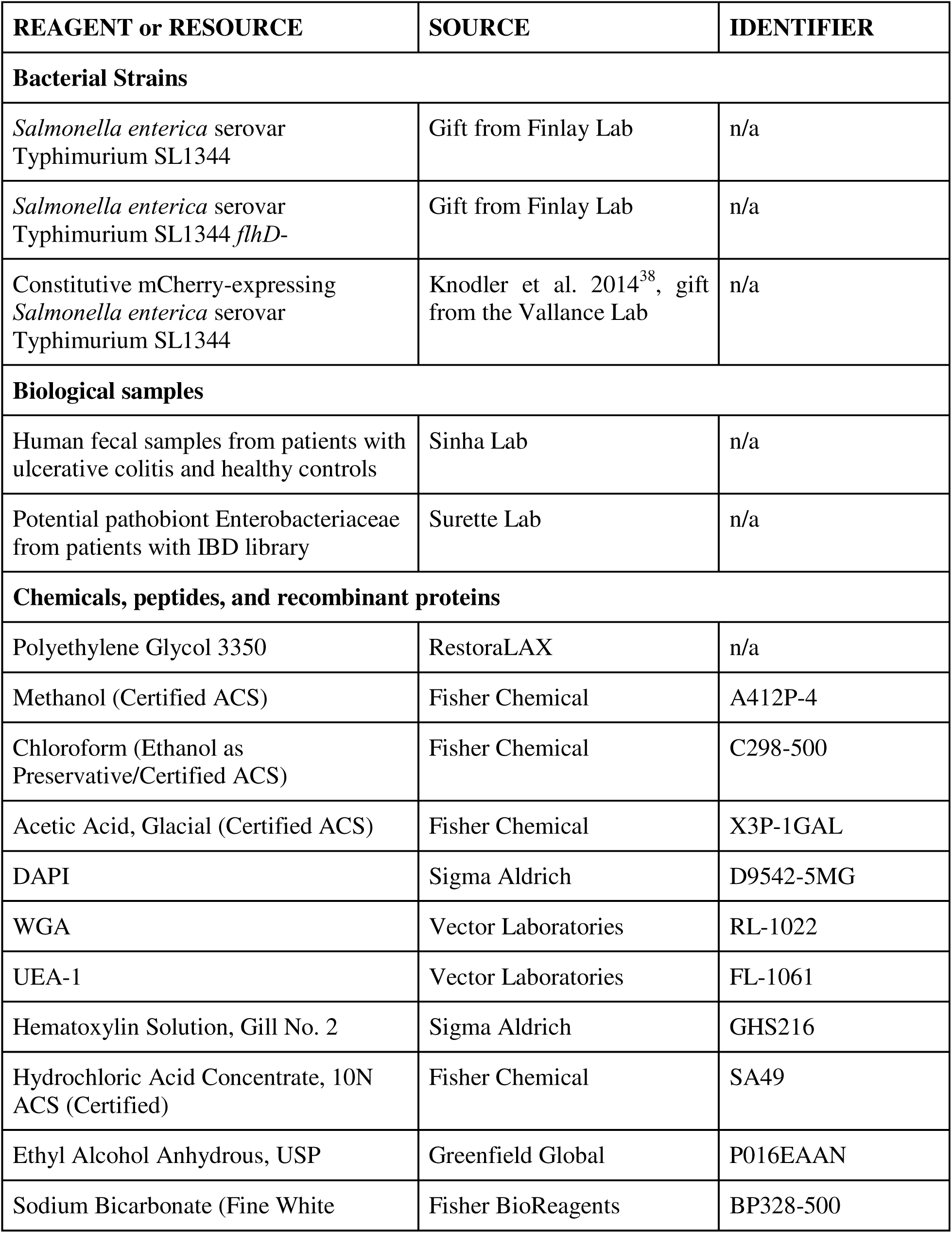
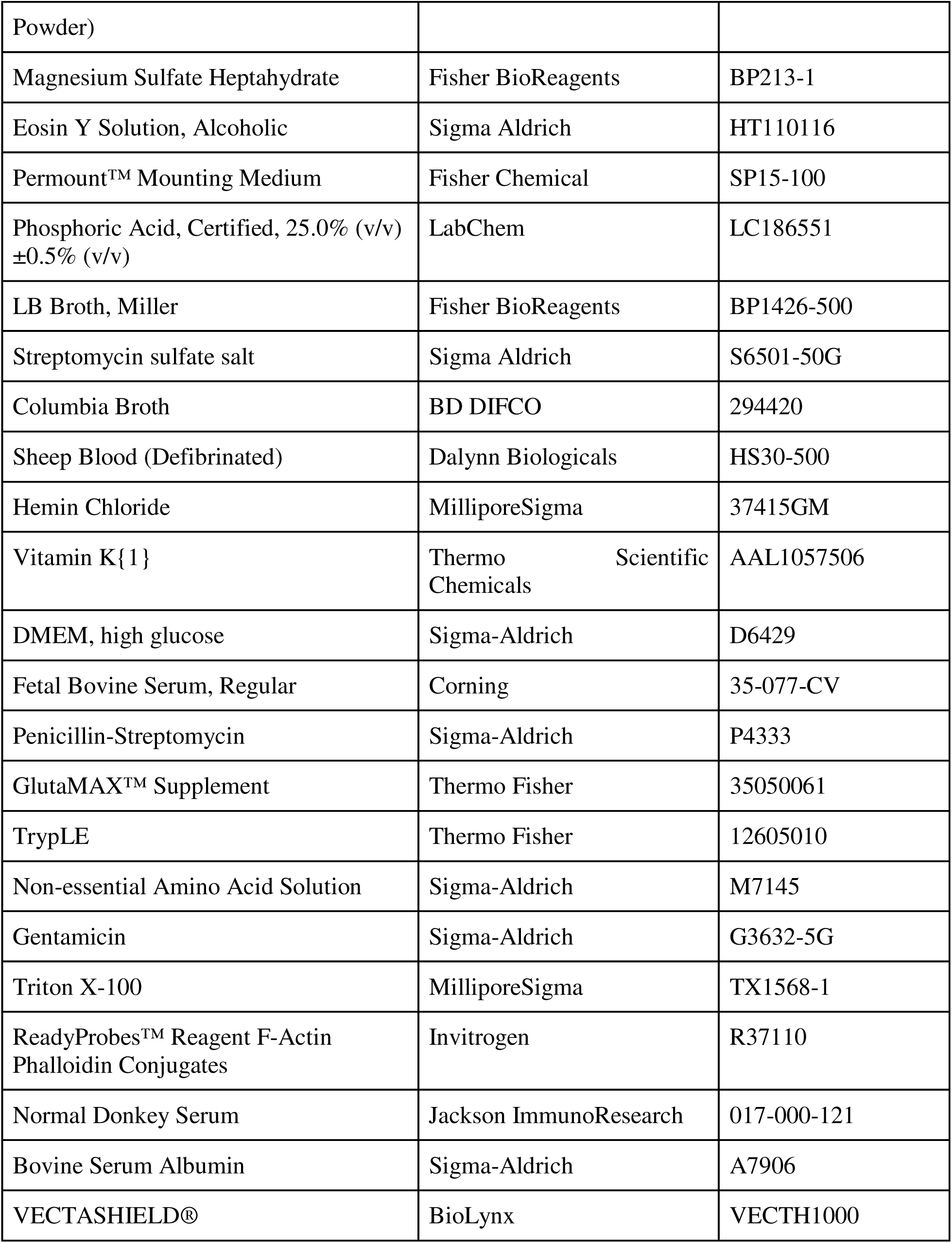
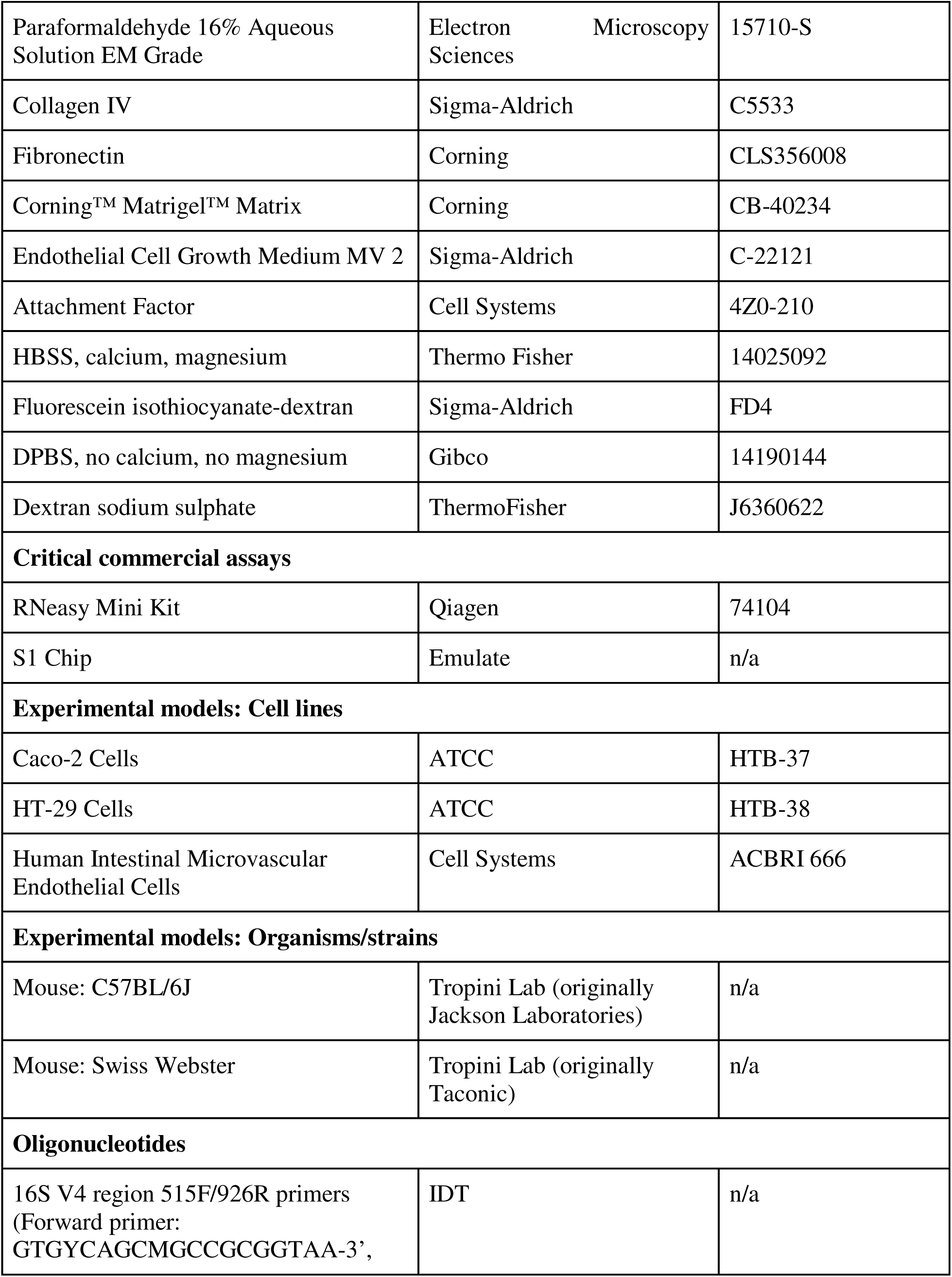
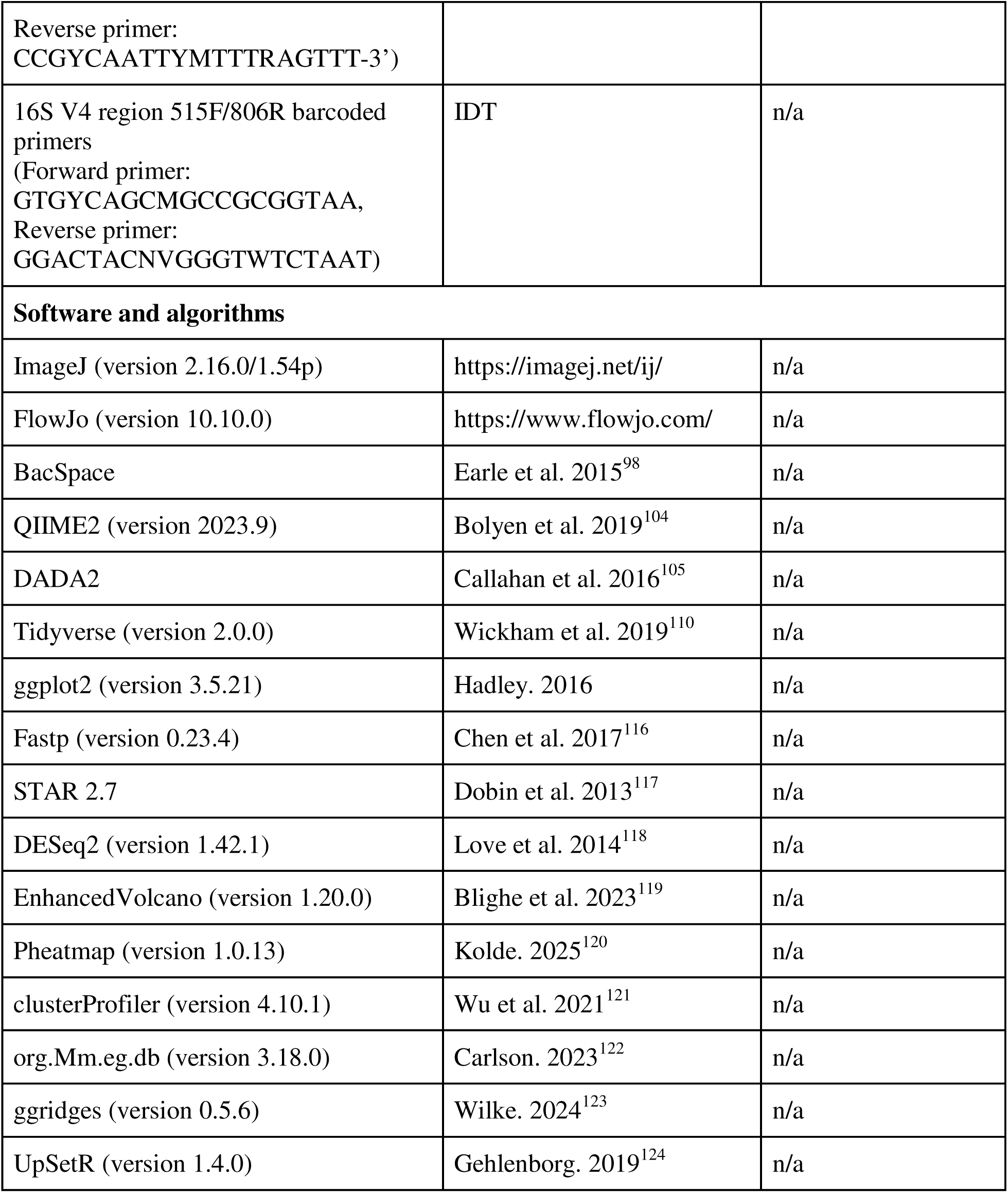

