## Supplementary figures and images for "In mouse and in vitro models, bowel preparation promotes pathogen colonization, translocation and exacerbation of inflammation"

### Supplemental Figure 1

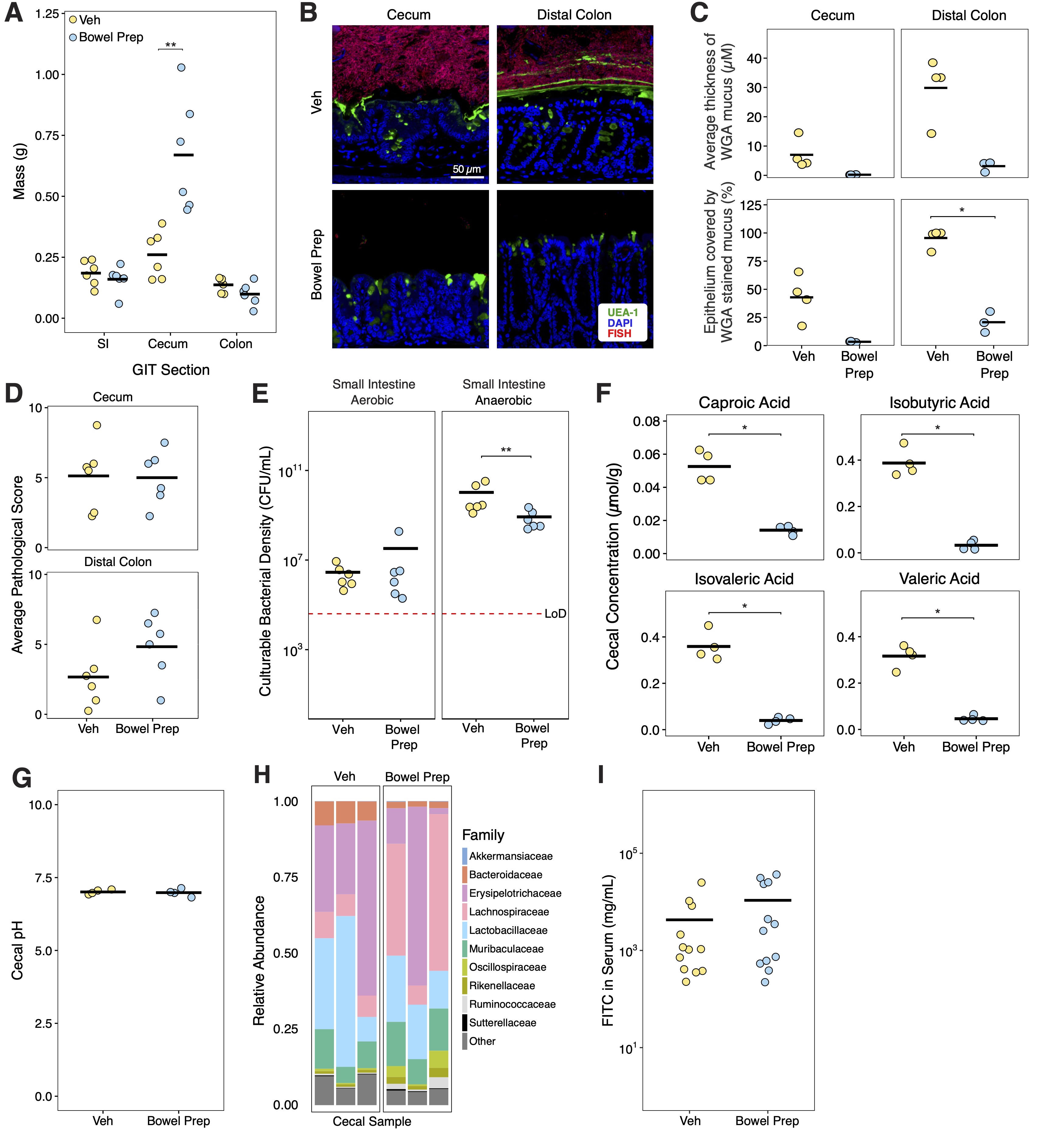

### Supplemental Figure 2

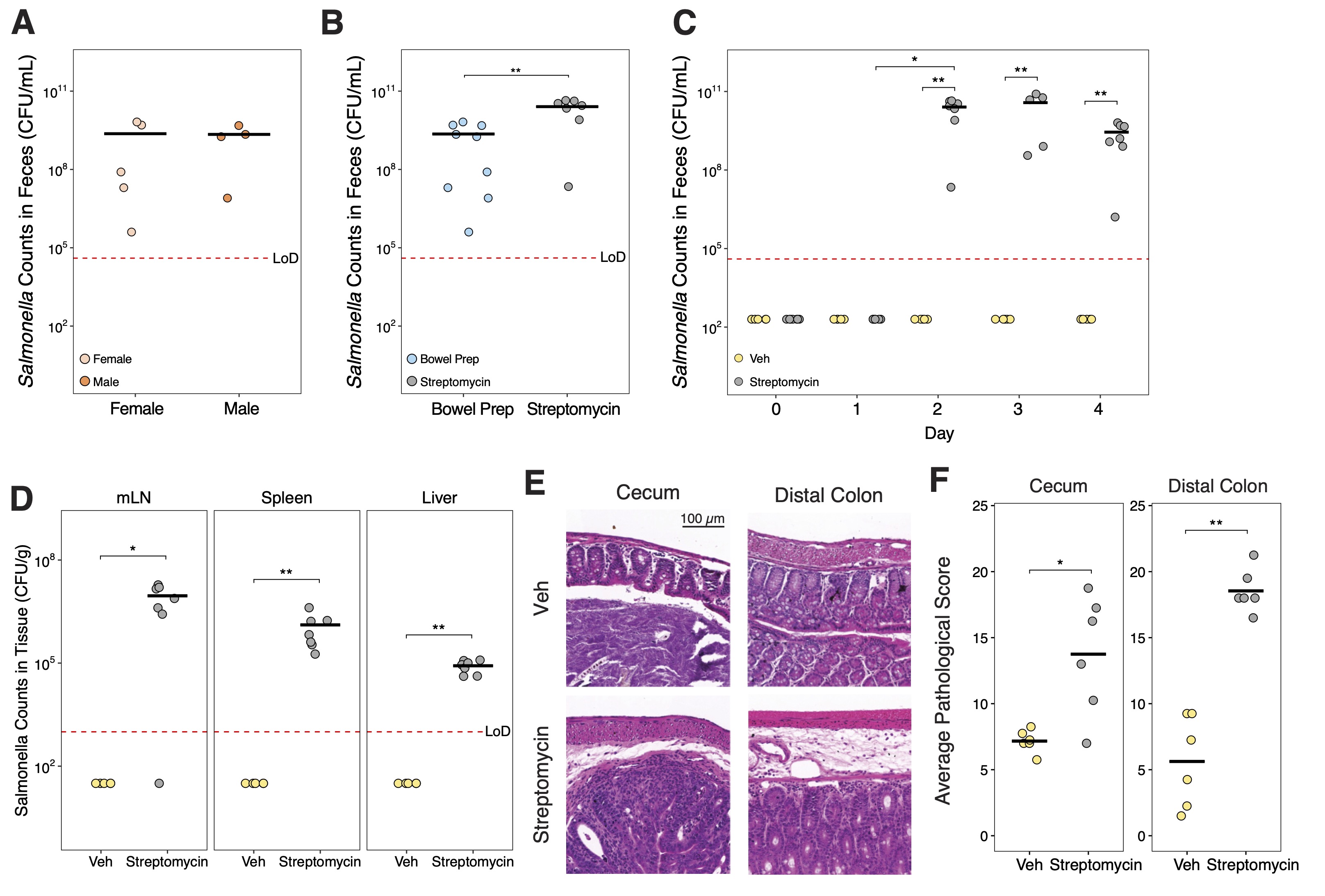

### Supplemental Figure 3

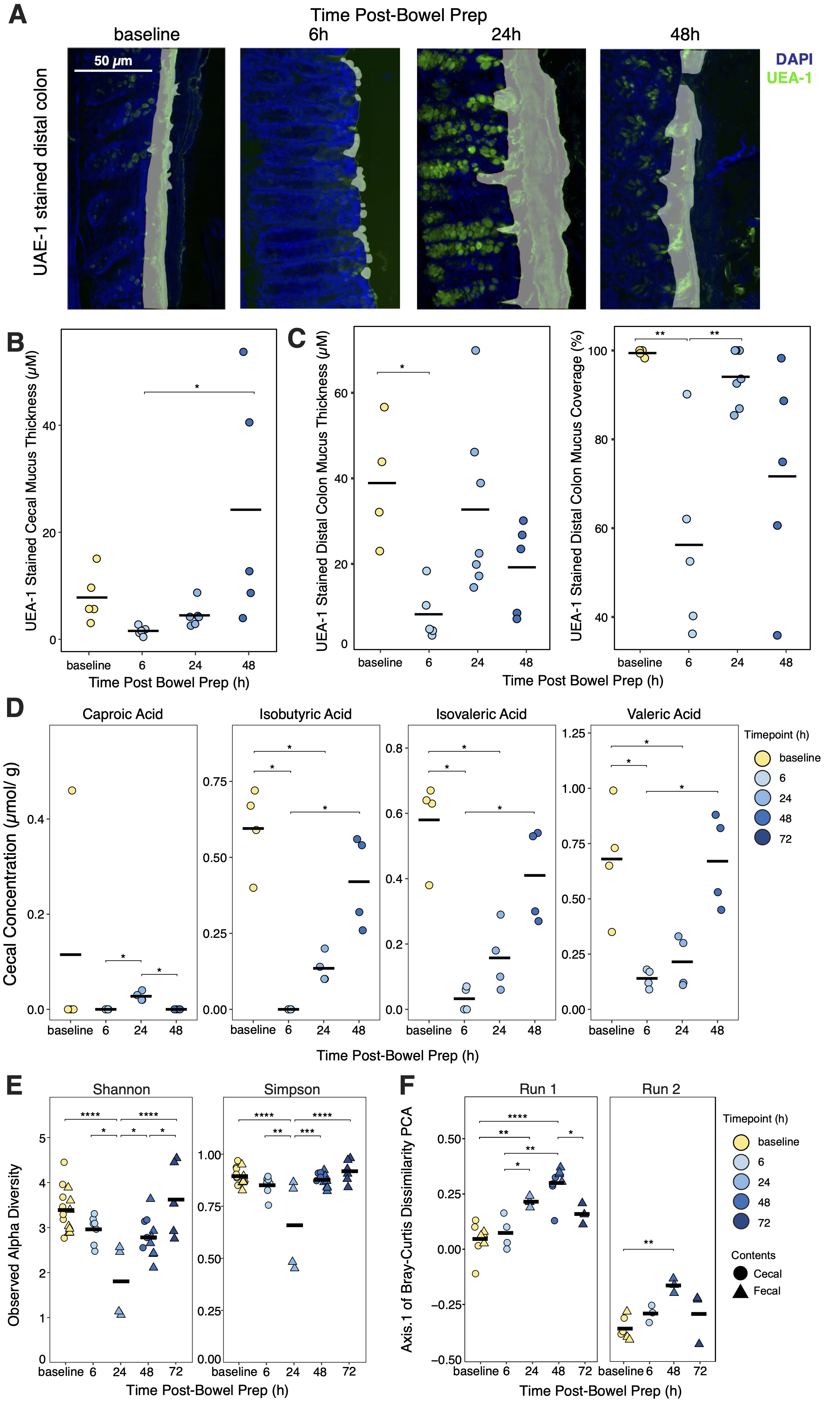

### Supplemental Figure 4

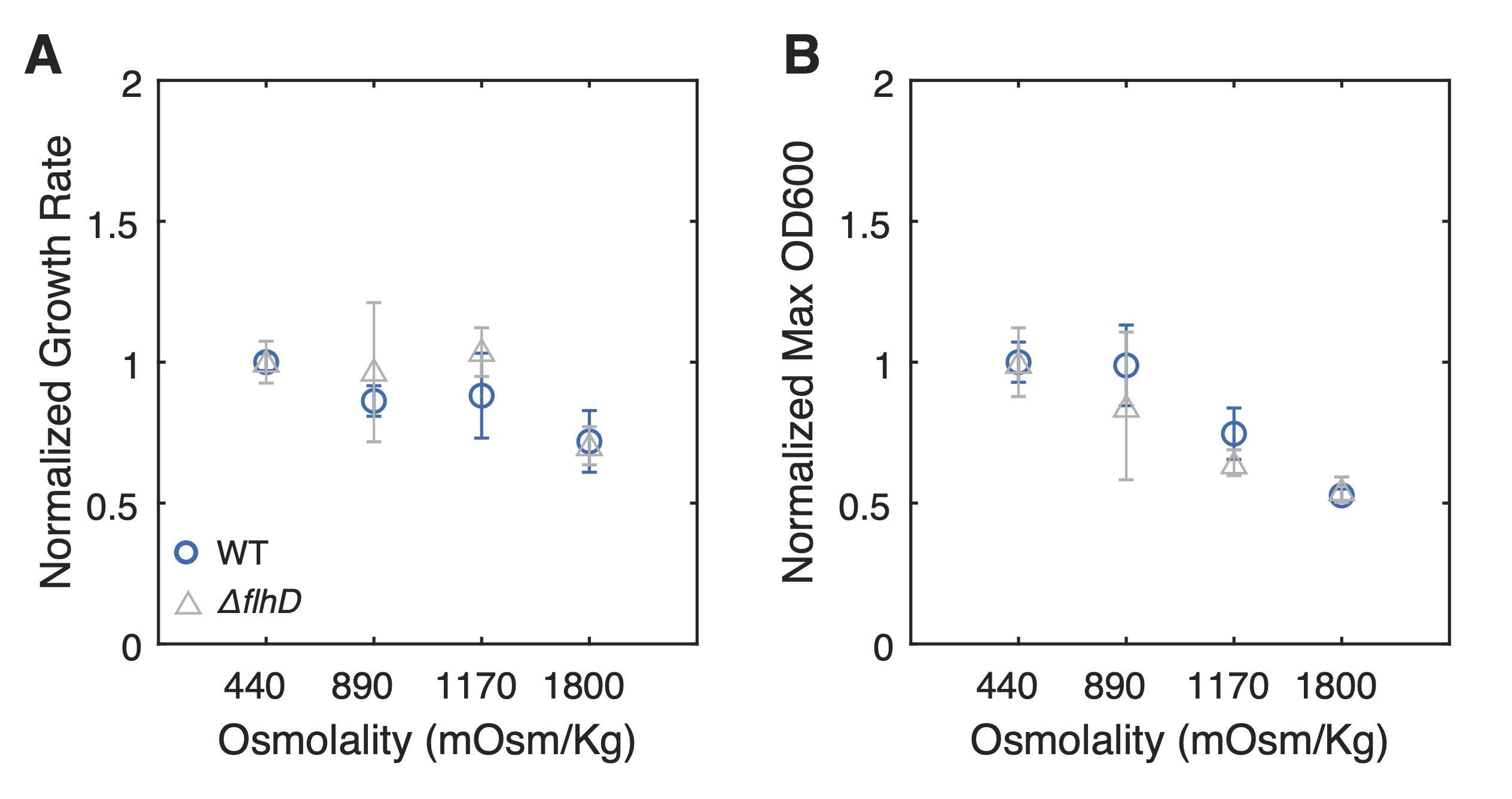

### Supplemental Figure 5

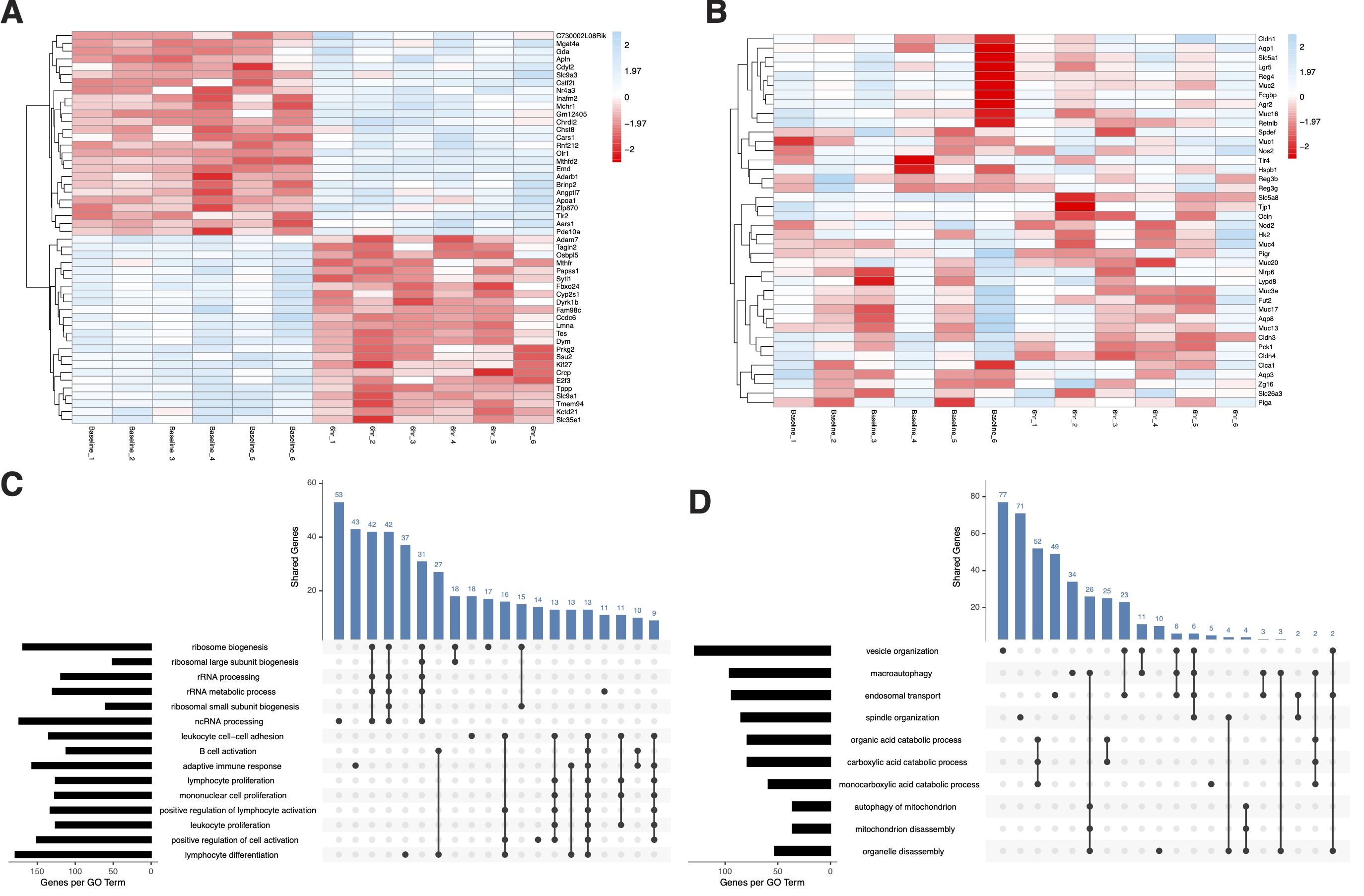

### Supplemental Figure 6

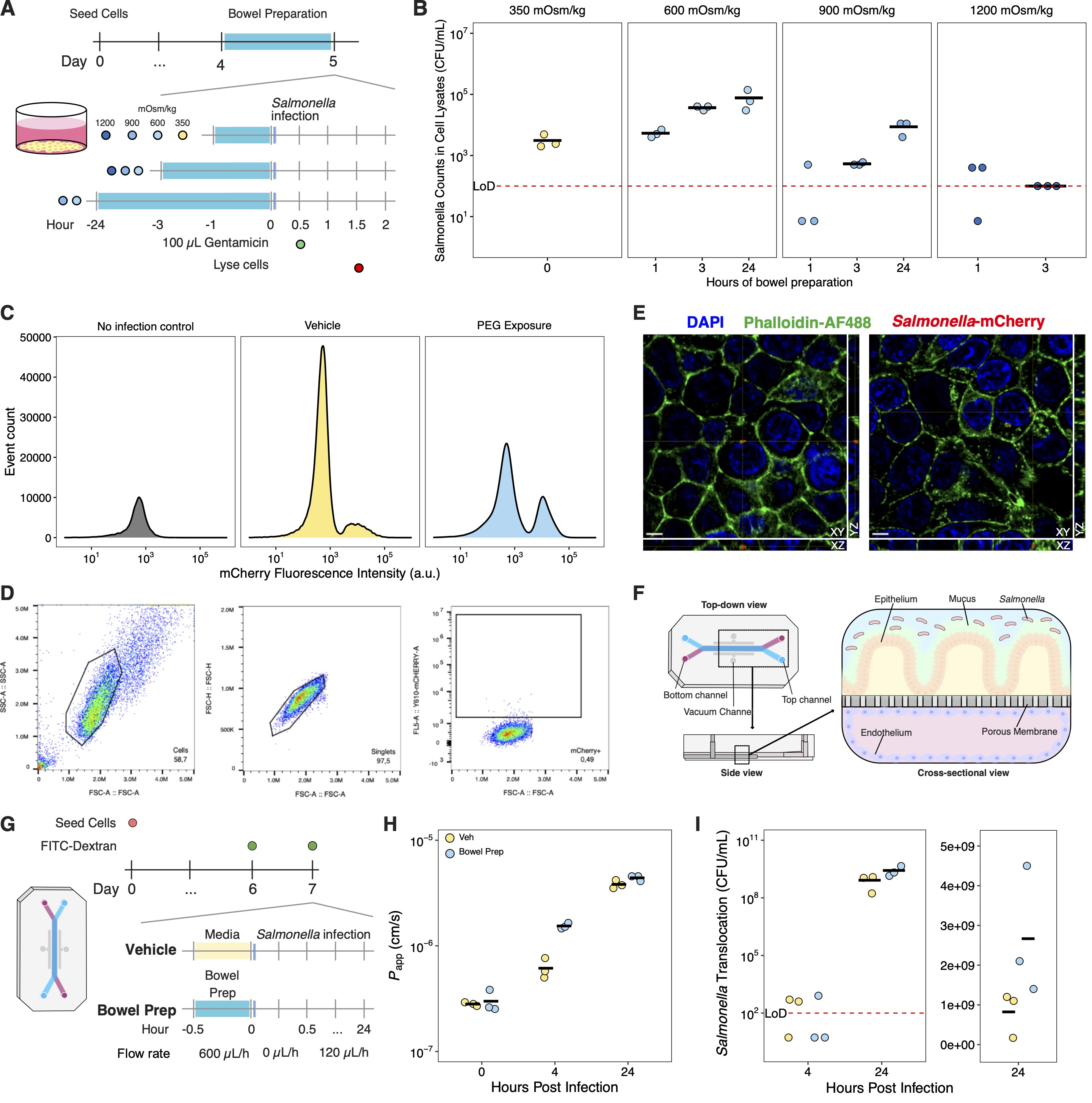

### Supplemental Figure 7

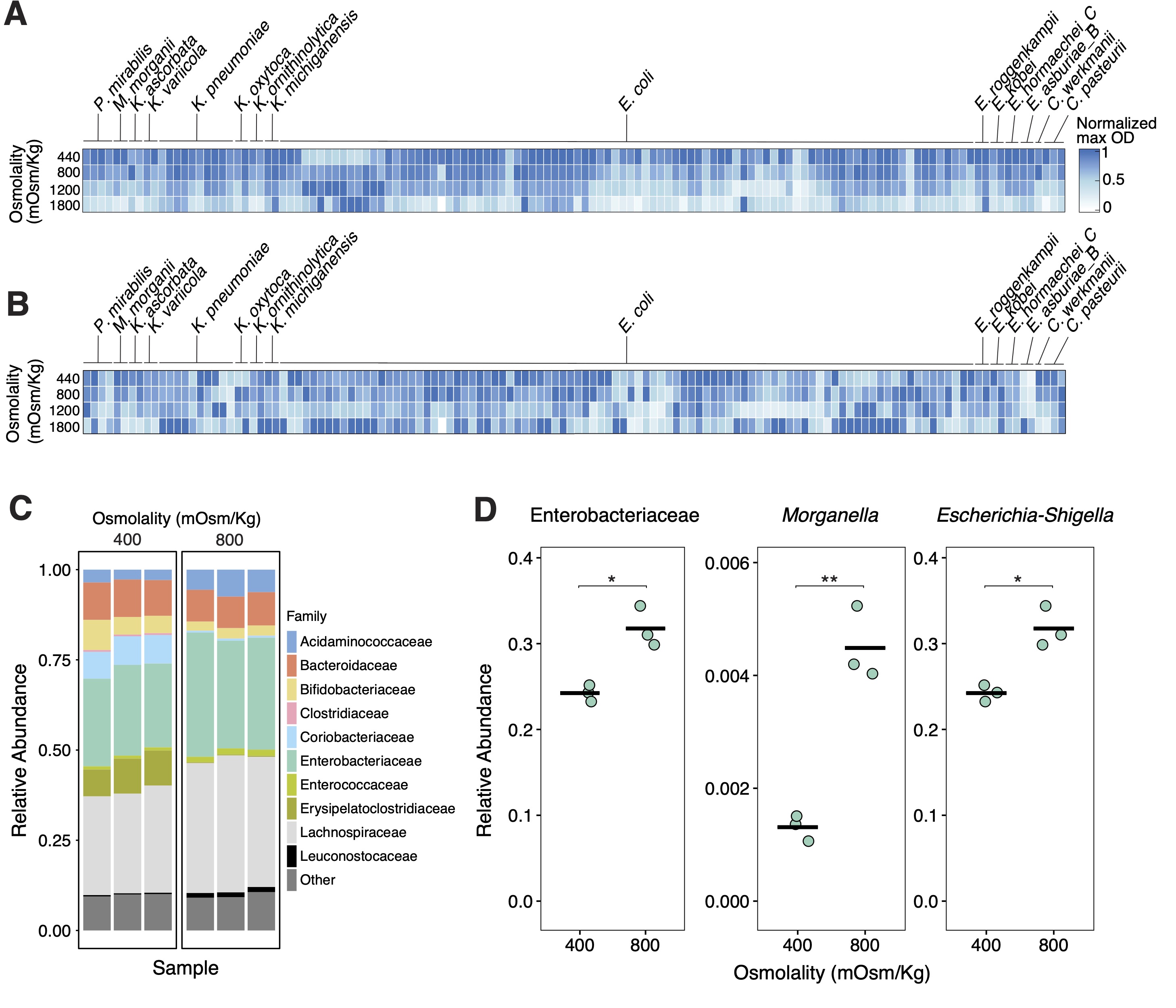

### Supplemental Figure 8

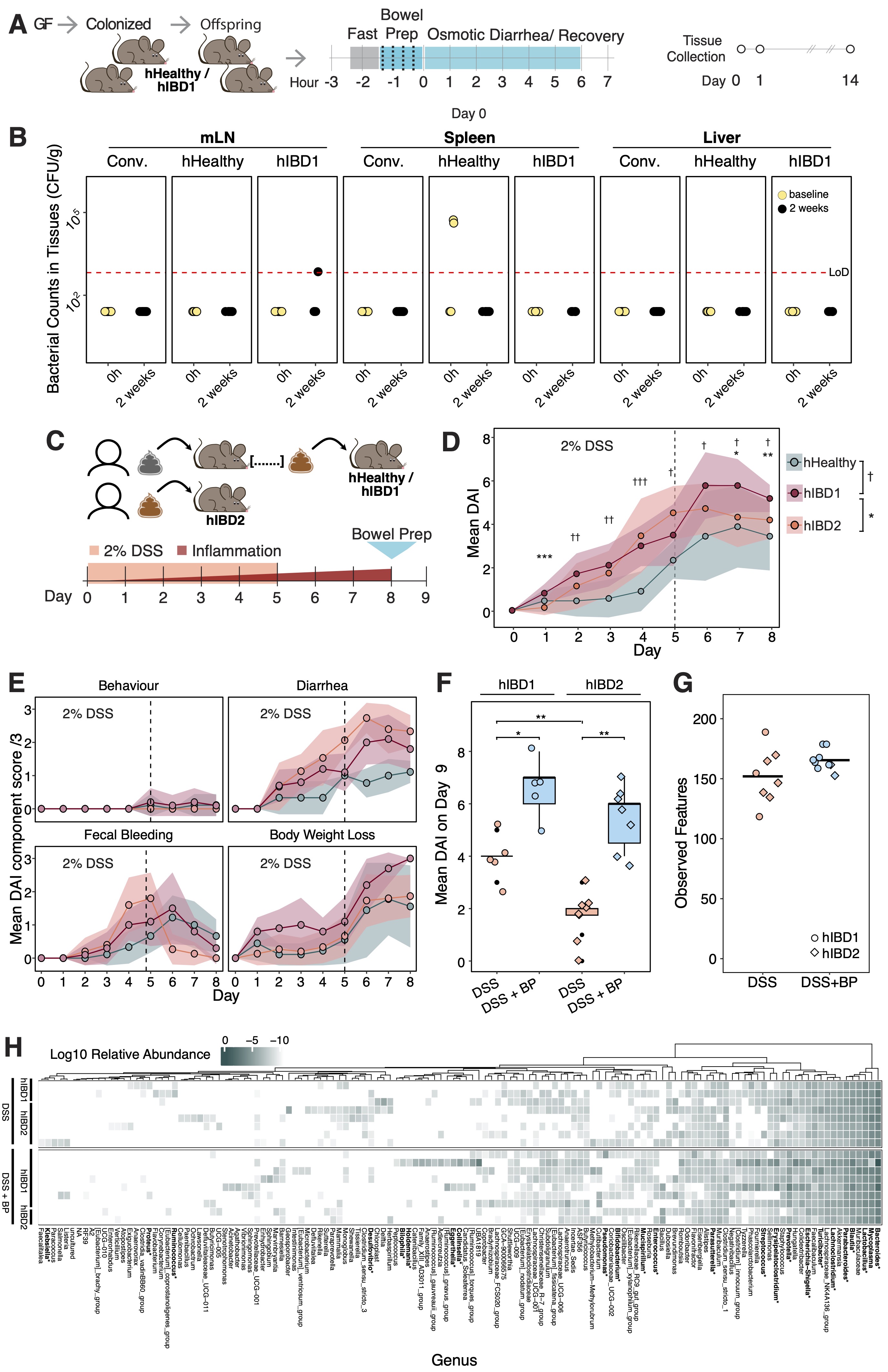
